# Evaluating the Combined Neurotoxicity of Amyloid Beta and Tau Oligomers in Alzheimer’s Disease: A Novel Cellular-Level Criterion

**DOI:** 10.1101/2024.11.03.621770

**Authors:** Andrey V. Kuznetsov

## Abstract

A criterion characterizing the combined neurotoxicity of amyloid beta and tau oligomers is suggested. A mathematical model for calculating the value of this criterion during senile plaque and neurofibrillary tangle (NFT) formation is proposed. Computations show that for physiologically relevant parameter values, the value of the criterion increases approximately linearly with time. Once neurofibrillary tangles begin forming in addition to senile plaques, there is an increase in the slope characterizing the rate at which the criterion increases. The critical value of the criterion at which a neuron dies is estimated. Unless the production rates of amyloid beta and tau monomers are very large, computations predict that for the accumulated toxicity to reach the critical value, the neural machinery responsible for the degradation of amyloid beta and tau monomers and aggregates must become dysfunctional. The value of the criterion after 20 years of the aggregation process is strongly influenced by the deposition rates of amyloid beta and tau oligomers into senile plaques and NFTs. This suggests that deposition of amyloid beta and tau oligomers into senile plaques and NFTs may reduce accumulated toxicity by sequestering more toxic oligomeric species into less toxic insoluble aggregates.

## 1. Introduction

Alzheimer’s disease (AD) is a neurodegenerative disorder marked by gradually worsening cognitive impairment. It is characterized by the buildup of amyloid-beta (Aβ) plaques outside neurons and neurofibrillary tangles (NFTs) composed of hyperphosphorylated tau protein within neurons, both of which are the hallmark features found in affected brain tissue [1-7].

Aβ plaques, also known as senile plaques, are the earliest pathological feature that develops in AD. The “amyloid cascade hypothesis” proposes that Aβ plaque aggregation is the primary process triggering neurotoxicity (hereafter toxicity) and dementia in AD [8,9]. Later research suggests that Aβ oligomers, rather than mature fibrils, are likely to be the most toxic Aβ species [10].

In AD, NFTs typically begin to develop about a decade after Aβ plaque formation starts [11]. NFTs, however, show a stronger correlation with cognitive decline compared to senile plaques [12]. Studies indicate that Aβ oligomers may seed the formation of NFTs [13]. Moreover, recent findings suggest that soluble tau oligomers are more harmful to neurons than NFTs [14]. The formation of NFTs may reduce accumulated toxicity by sequestering oligomeric tau species into less toxic insoluble aggregates [15].

This paper introduces a new criterion for characterizing the toxicity resulting from the combined effects of Aβ and tau oligomers. It builds on previous research into the role of Aβ oligomers in AD [16] and α-synuclein oligomers in Parkinson’s disease [17].

## 2. Materials and models

### 2.1. Equations simulating aggregation of Aβ peptide

The Finke−Watzky (F-W) model, described in refs. [18-20], is used to simulate the aggregation of Aβ peptides, representing aggregate growth as a two-step process involving nucleation and autocatalysis. During the nucleation step, new Aβ aggregates form continuously, typically at a slow rate. In the autocatalytic step, existing Aβ aggregates catalyze their own production. The autocatalytic process is generally much faster than nucleation. These two pseudo-elementary reaction steps are described as follows:

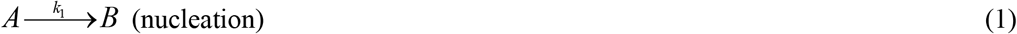

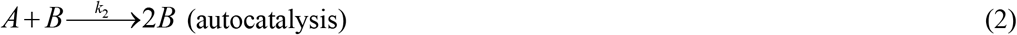

where *A* denotes monomeric Aβ peptides, while *B* represents free Aβ aggregates, such as oligomers and protofibrils, which have not yet been incorporated into senile plaques. From this point forward, the terms “free Aβ aggregates” and “Aβ oligomers” are used interchangeably, although “free Aβ aggregates” may encompass dimers, oligomers, and protofibrils [21]. The rate constants *k*_1_ and *k*_2_ define the nucleation and autocatalytic growth rates, respectively [18]. Primary nucleation, as expressed in Eq. (1), involves only monomers, whereas secondary nucleation, described in Eq. (2), involves both monomers and pre-existing free aggregates of the same peptide [22]. Analytical solutions for the F-W model, addressing the conversion of a fixed amount of monomers into polymers, are well-established in the literature [23]. However, this study modifies the F-W model to simulate a different context, specifically the scenario in which monomers are continuously produced.

Table 1 summarizes the dependent variables used in the model, while Table 2 summarizes the model parameters. Ref. [24] introduced a dimensionless parameter, *ξ*_*Aβ*_, representing the ratio of the variation in the Aβ monomer concentration across the control volume (CV) to the average Aβ monomer concentration within the CV at time *t*_*f*_ :

**Table 1.**
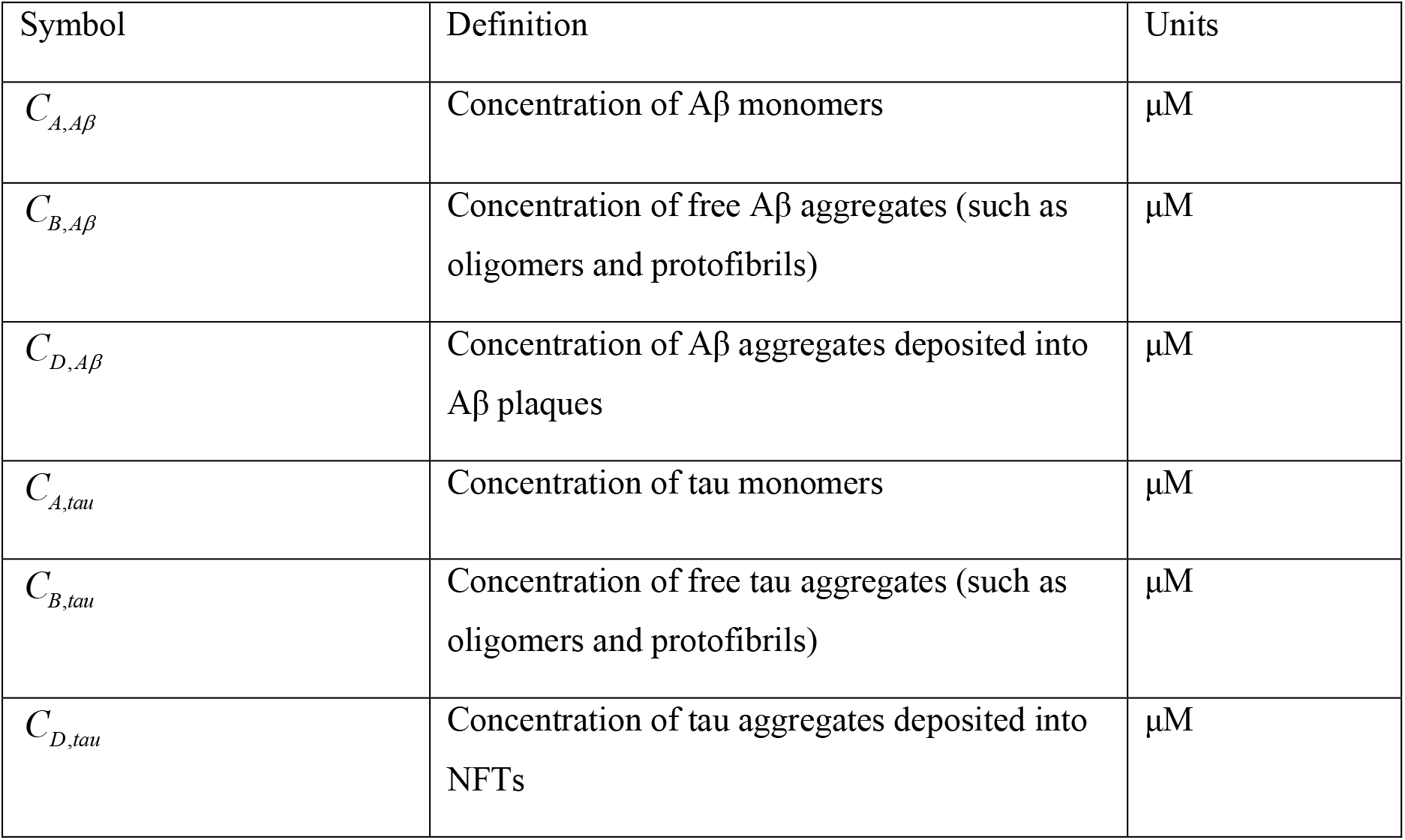
Dependent variables utilized in the model.

| Symbol | Definition | Units |
| --- | --- | --- |
| $C_{A,A\beta}$ | Concentration of Aβ monomers | $\mu\text{M}$ |
| $C_{B,A\beta}$ | Concentration of free Aβ aggregates (such as oligomers and protofibrils) | $\mu\text{M}$ |
| $C_{D,A\beta}$ | Concentration of Aβ aggregates deposited into Aβ plaques | $\mu\text{M}$ |
| $C_{A,\tau}$ | Concentration of tau monomers | $\mu\text{M}$ |
| $C_{B,\tau}$ | Concentration of free tau aggregates (such as oligomers and protofibrils) | $\mu\text{M}$ |
| $C_{D,\tau}$ | Concentration of tau aggregates deposited into NFTs | $\mu\text{M}$ |

**Table 2.**
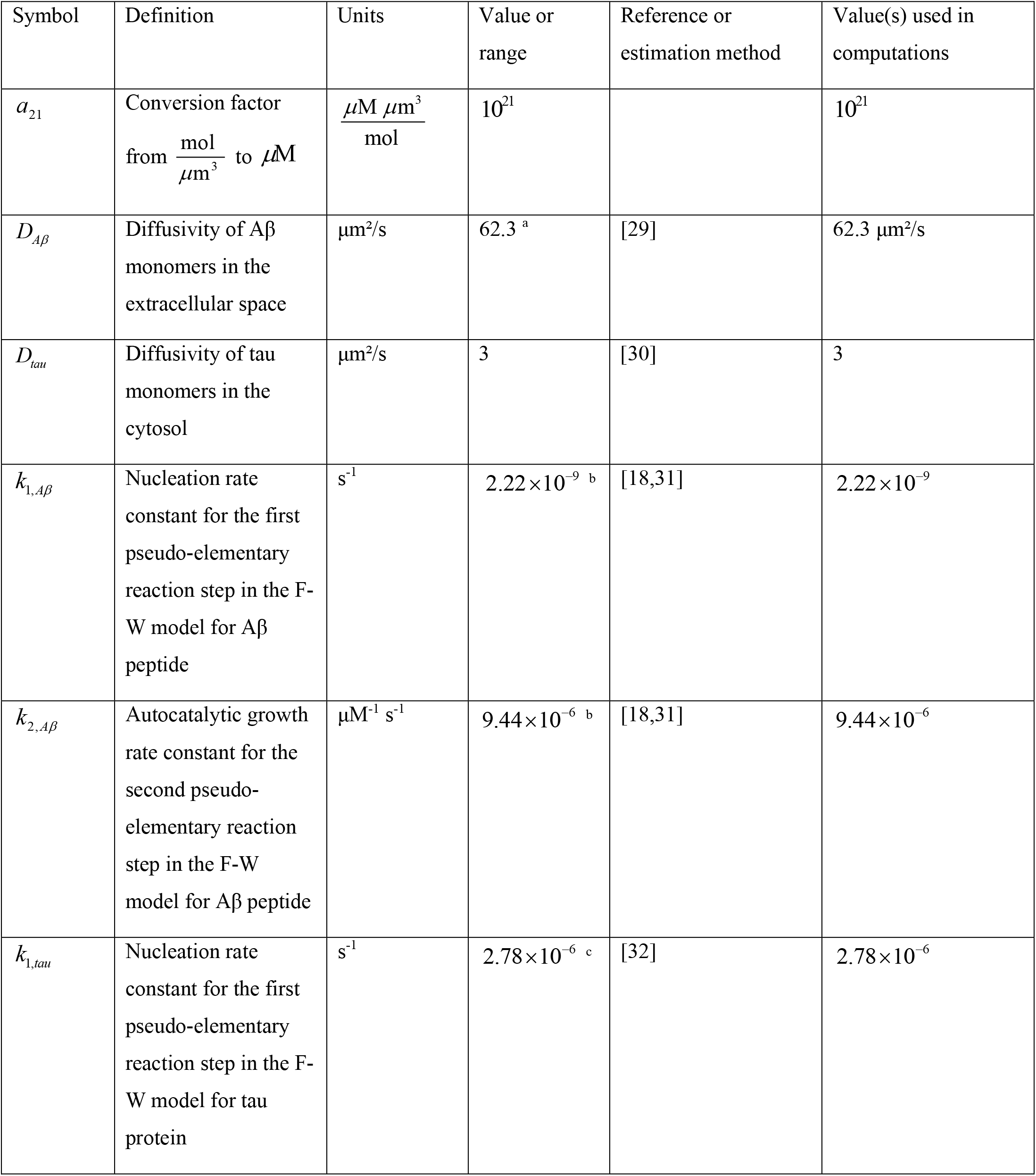
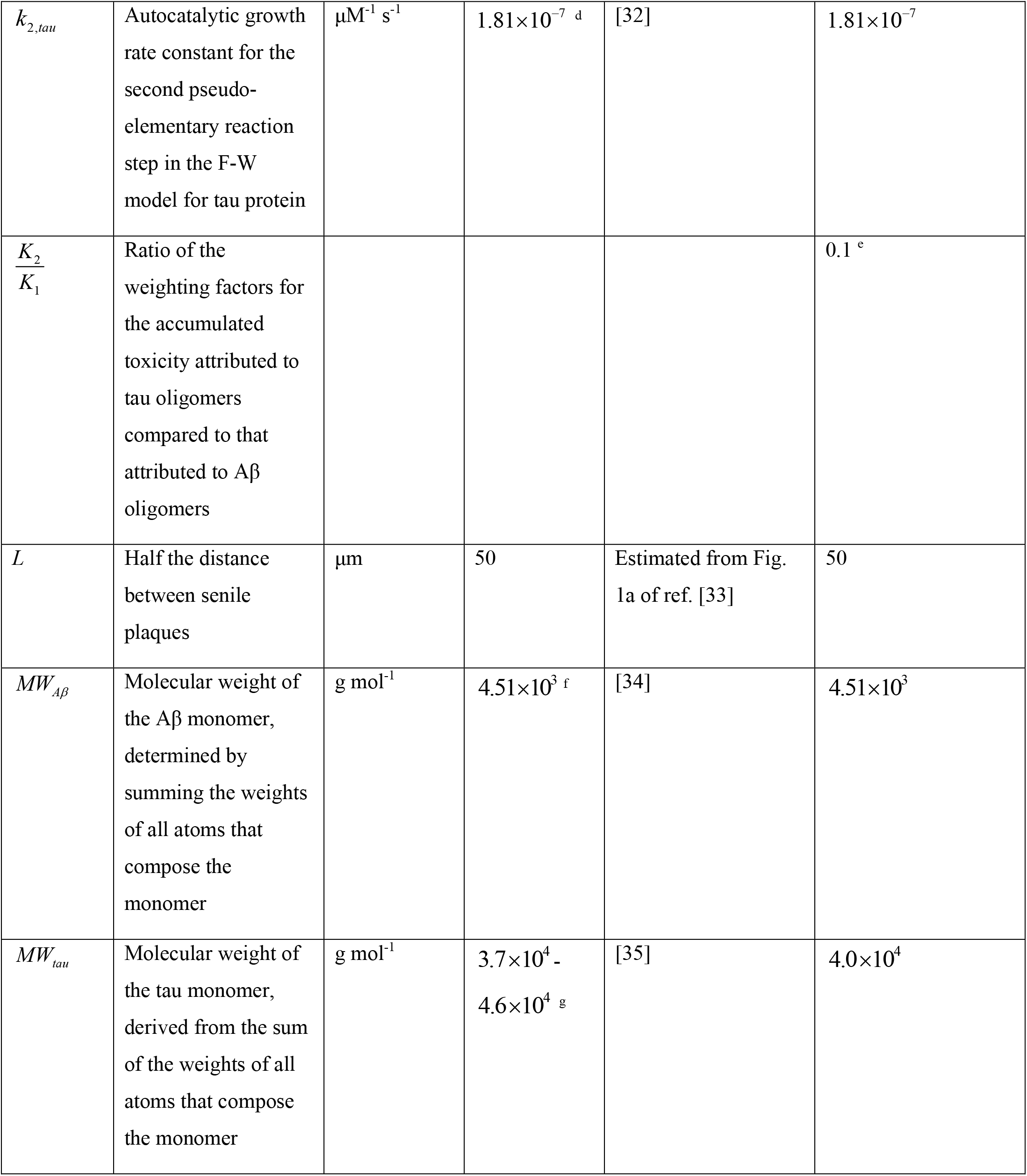
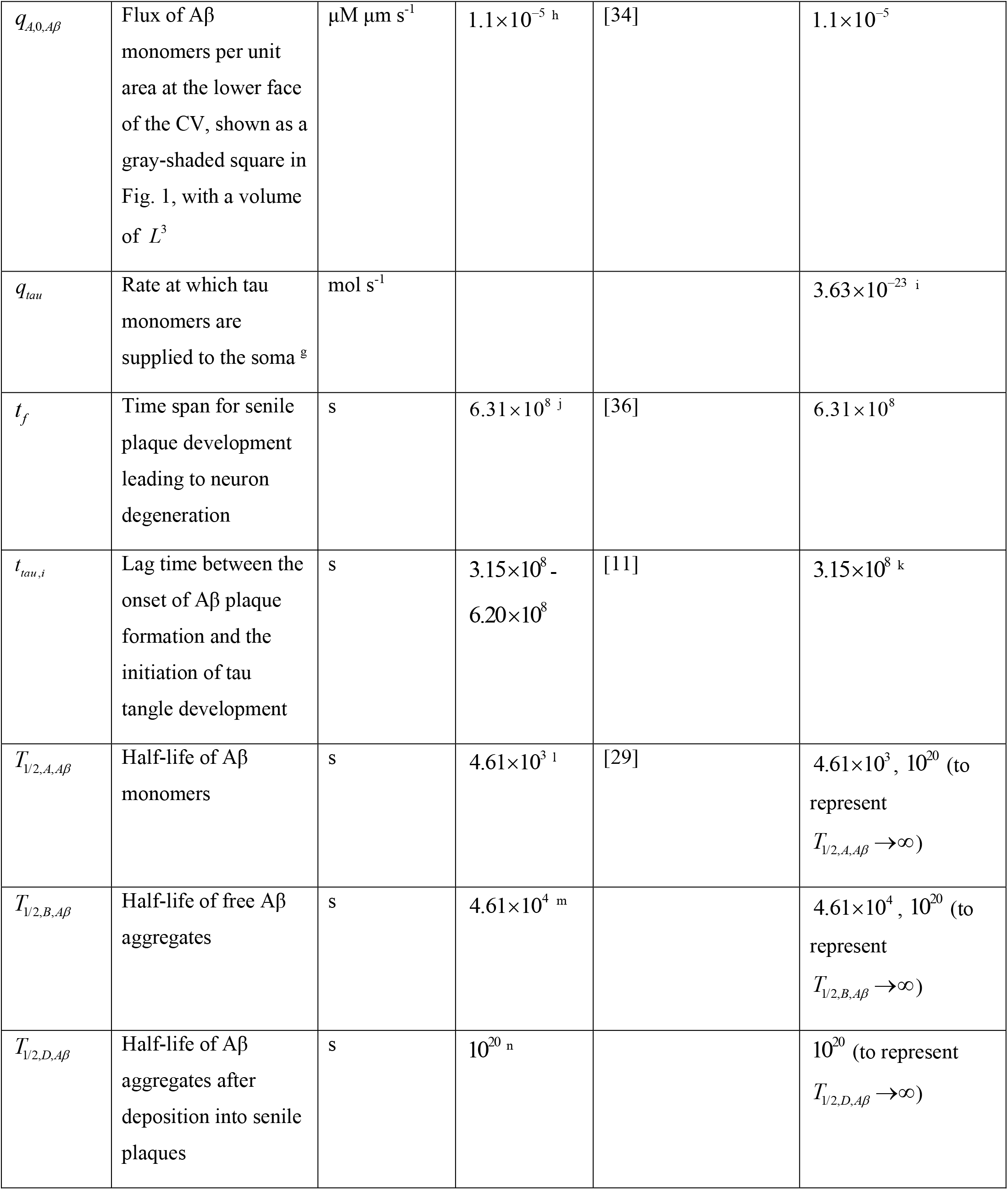
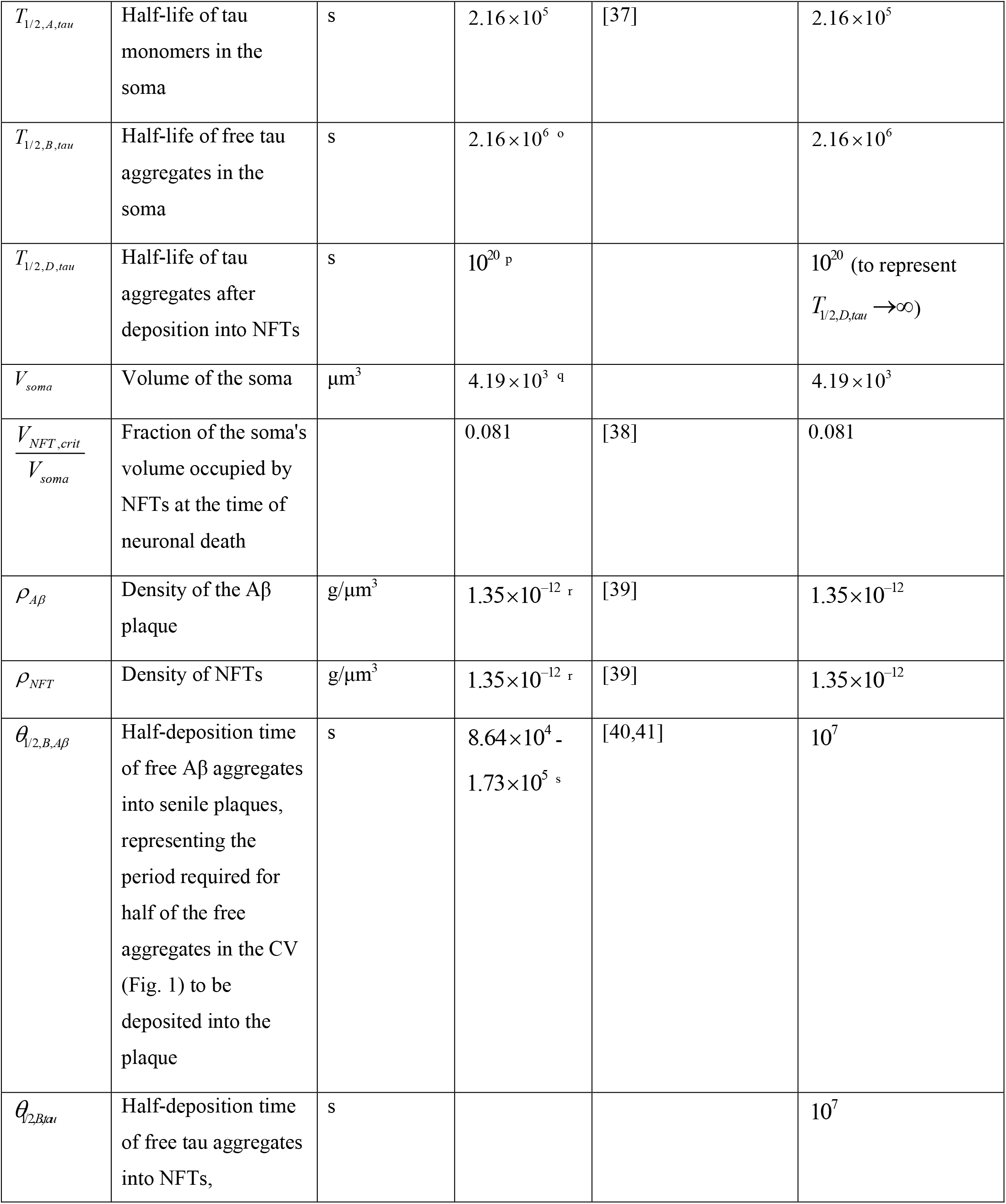
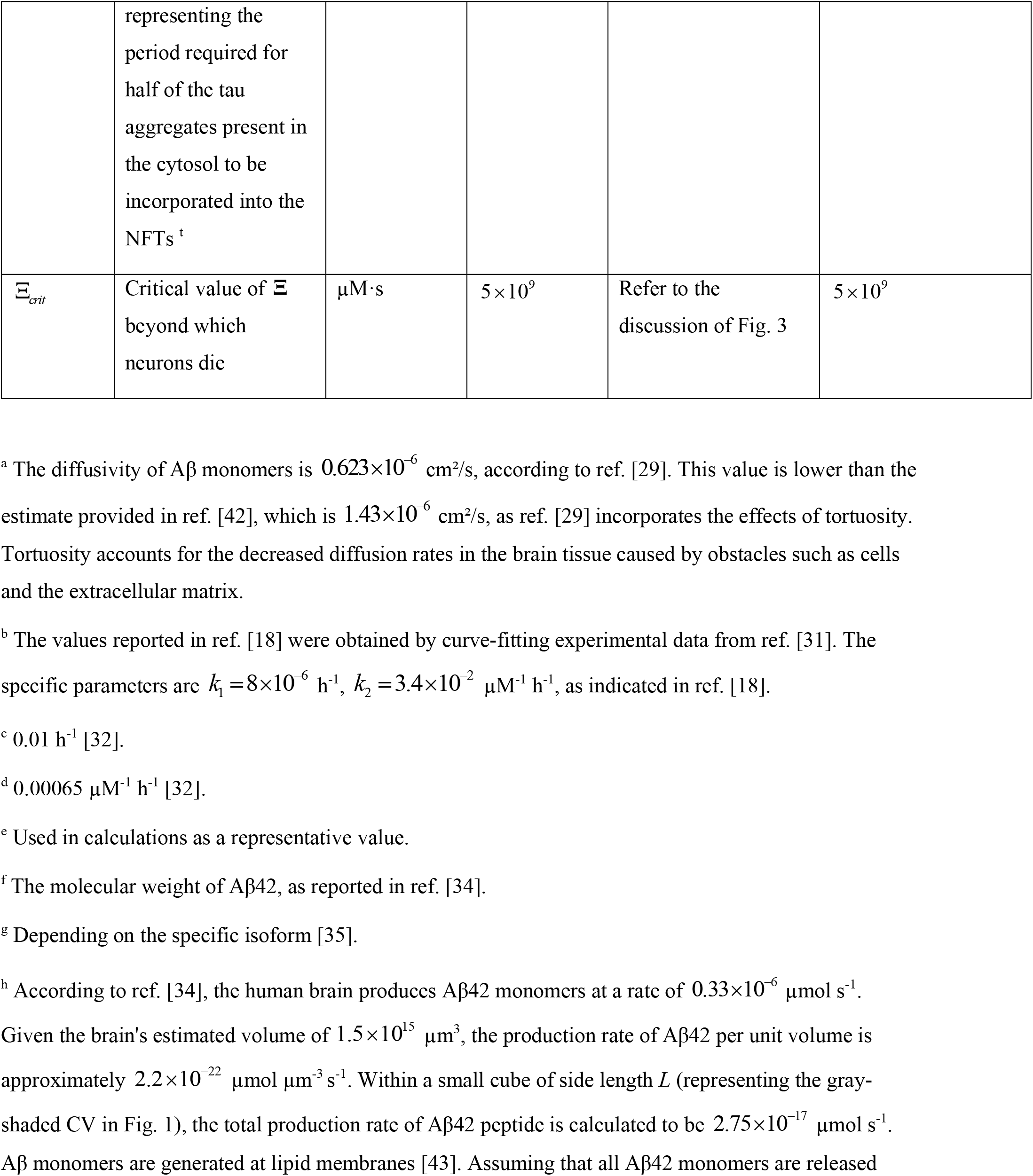
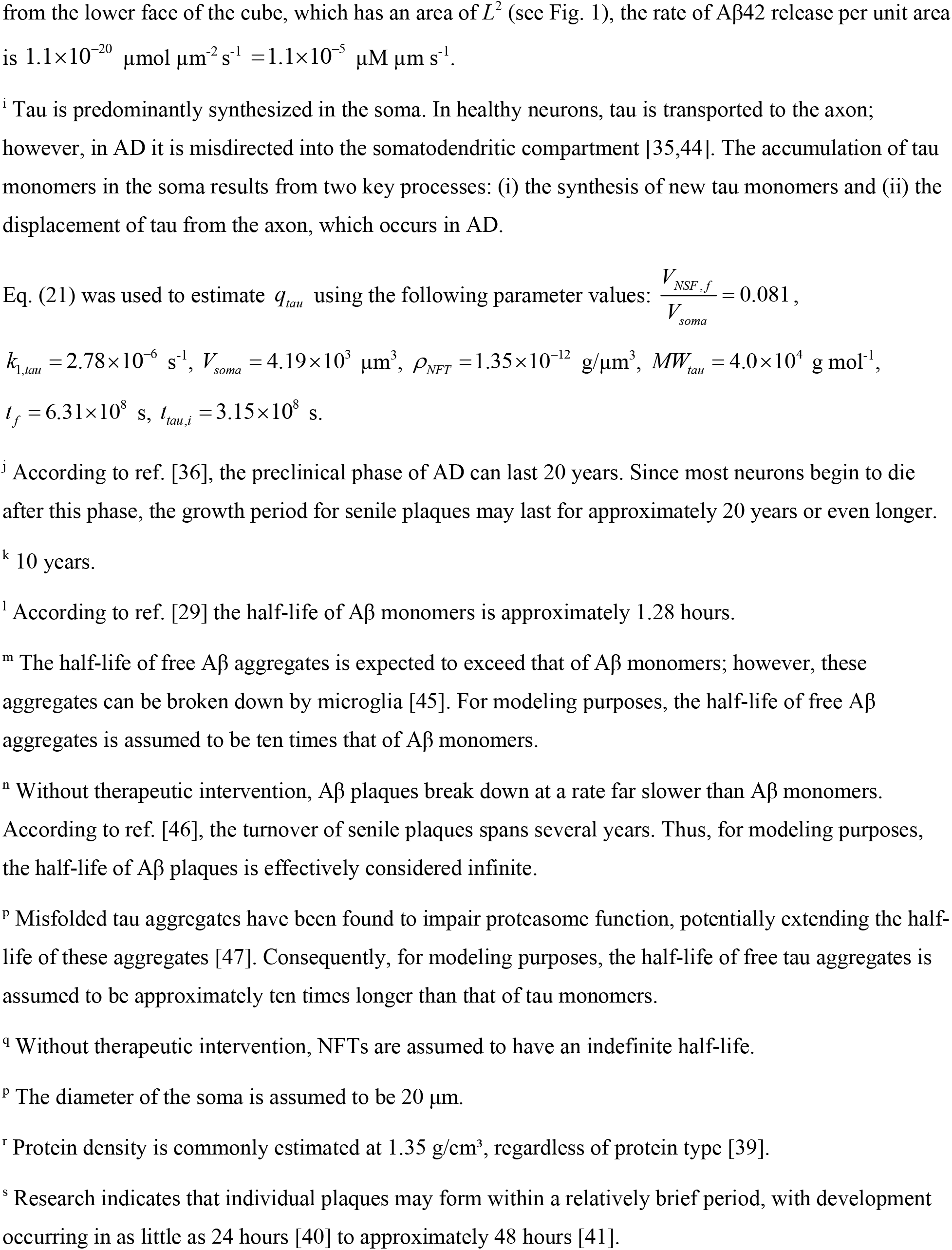
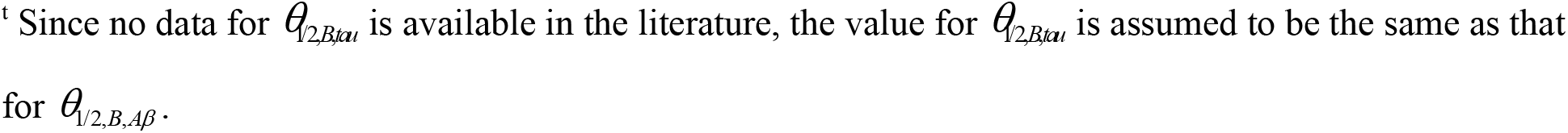
Model parameters.

| Symbol | Definition | Units | Value or range | Reference or estimation method | Value(s) used in computations |
| --- | --- | --- | --- | --- | --- |
| $a_{21}$ | Conversion factor from $\frac{\text{mol}}{\mu\text{m}^3}$ to $\mu\text{M}$ | $\frac{\mu\text{M } \mu\text{m}^3}{\text{mol}}$ | $10^{21}$ | | $10^{21}$ |
| $D_{A\beta}$ | Diffusivity of Aβ monomers in the extracellular space | $\mu\text{m}^2/\text{s}$ | 62.3 <sup>a</sup> | [29] | 62.3 $\mu\text{m}^2/\text{s}$ |
| $D_{\text{tau}}$ | Diffusivity of tau monomers in the cytosol | $\mu\text{m}^2/\text{s}$ | 3 | [30] | 3 |
| $k_{1,A\beta}$ | Nucleation rate constant for the first pseudo-elementary reaction step in the F-W model for Aβ peptide | $\text{s}^{-1}$ | $2.22 \times 10^{-9}$ <sup>b</sup> | [18,31] | $2.22 \times 10^{-9}$ |
| $k_{2,A\beta}$ | Autocatalytic growth rate constant for the second pseudo-elementary reaction step in the F-W model for Aβ peptide | $\mu\text{M}^{-1} \text{s}^{-1}$ | $9.44 \times 10^{-6}$ <sup>b</sup> | [18,31] | $9.44 \times 10^{-6}$ |
| $k_{1,\text{tau}}$ | Nucleation rate constant for the first pseudo-elementary reaction step in the F-W model for tau protein | $\text{s}^{-1}$ | $2.78 \times 10^{-6}$ <sup>c</sup> | [32] | $2.78 \times 10^{-6}$ |
| $k_{2,\tau}$ | Autocatalytic growth rate constant for the second pseudo-elementary reaction step in the F-W model for tau protein | $\mu\text{M}^{-1} \text{s}^{-1}$ | $1.81 \times 10^{-7} \text{ }^{\text{d}}$ | [32] | $1.81 \times 10^{-7}$ |
| $\frac{K_2}{K_1}$ | Ratio of the weighting factors for the accumulated toxicity attributed to tau oligomers compared to that attributed to Aβ oligomers | | | | $0.1 \text{ }^{\text{e}}$ |
| $L$ | Half the distance between senile plaques | $\mu\text{m}$ | 50 | Estimated from Fig. 1a of ref. [33] | 50 |
| $MW_{A\beta}$ | Molecular weight of the Aβ monomer, determined by summing the weights of all atoms that compose the monomer | $\text{g mol}^{-1}$ | $4.51 \times 10^3 \text{ }^{\text{f}}$ | [34] | $4.51 \times 10^3$ |
| $MW_{\tau}$ | Molecular weight of the tau monomer, derived from the sum of the weights of all atoms that compose the monomer | $\text{g mol}^{-1}$ | $3.7 \times 10^4 - 4.6 \times 10^4 \text{ }^{\text{g}}$ | [35] | $4.0 \times 10^4$ |
| $q_{A,0,A\beta}$ | Flux of Aβ monomers per unit area at the lower face of the CV, shown as a gray-shaded square in Fig. 1, with a volume of $L^3$ | $\mu\text{M } \mu\text{m s}^{-1}$ | $1.1 \times 10^{-5}$ <sup>h</sup> | [34] | $1.1 \times 10^{-5}$ |
| $q_{\text{tau}}$ | Rate at which tau monomers are supplied to the soma <sup>g</sup> | $\text{mol s}^{-1}$ | | | $3.63 \times 10^{-23}$ <sup>i</sup> |
| $t_f$ | Time span for senile plaque development leading to neuron degeneration | s | $6.31 \times 10^8$ <sup>j</sup> | [36] | $6.31 \times 10^8$ |
| $t_{\text{tau},i}$ | Lag time between the onset of Aβ plaque formation and the initiation of tau tangle development | s | $3.15 \times 10^8$ - $6.20 \times 10^8$ | [11] | $3.15 \times 10^8$ <sup>k</sup> |
| $T_{1/2,A,A\beta}$ | Half-life of Aβ monomers | s | $4.61 \times 10^3$ <sup>l</sup> | [29] | $4.61 \times 10^3$, $10^{20}$ (to represent $T_{1/2,A,A\beta} \rightarrow \infty$) |
| $T_{1/2,B,A\beta}$ | Half-life of free Aβ aggregates | s | $4.61 \times 10^4$ <sup>m</sup> | | $4.61 \times 10^4$, $10^{20}$ (to represent $T_{1/2,B,A\beta} \rightarrow \infty$) |
| $T_{1/2,D,A\beta}$ | Half-life of Aβ aggregates after deposition into senile plaques | s | $10^{20}$ <sup>n</sup> | | $10^{20}$ (to represent $T_{1/2,D,A\beta} \rightarrow \infty$) |
| $T_{1/2,A,\tau}$ | Half-life of tau monomers in the soma | s | $2.16 \times 10^5$ | [37] | $2.16 \times 10^5$ |
| $T_{1/2,B,\tau}$ | Half-life of free tau aggregates in the soma | s | $2.16 \times 10^6$ ° | | $2.16 \times 10^6$ |
| $T_{1/2,D,\tau}$ | Half-life of tau aggregates after deposition into NFTs | s | $10^{20}$ p | | $10^{20}$ (to represent $T_{1/2,D,\tau} \rightarrow \infty$) |
| $V_{soma}$ | Volume of the soma | $\mu\text{m}^3$ | $4.19 \times 10^3$ q | | $4.19 \times 10^3$ |
| $\frac{V_{NFT,crit}}{V_{soma}}$ | Fraction of the soma's volume occupied by NFTs at the time of neuronal death | | 0.081 | [38] | 0.081 |
| $\rho_{A\beta}$ | Density of the Aβ plaque | $\text{g}/\mu\text{m}^3$ | $1.35 \times 10^{-12}$ r | [39] | $1.35 \times 10^{-12}$ |
| $\rho_{NFT}$ | Density of NFTs | $\text{g}/\mu\text{m}^3$ | $1.35 \times 10^{-12}$ r | [39] | $1.35 \times 10^{-12}$ |
| $\theta_{1/2,B,A\beta}$ | Half-deposition time of free Aβ aggregates into senile plaques, representing the period required for half of the free aggregates in the CV (Fig. 1) to be deposited into the plaque | s | $8.64 \times 10^4$ -<br>$1.73 \times 10^5$ s | [40,41] | $10^7$ |
| $\theta_{1/2,B,\tau}$ | Half-deposition time of free tau aggregates into NFTs, representing the period required for half of the tau aggregates present in the cytosol to be incorporated into the NFTs<sup>t</sup> | s | | | $10^7$ |
| $\Xi_{crit}$ | Critical value of $\Xi$ beyond which neurons die | $\mu\text{M}\cdot\text{s}$ | $5 \times 10^9$ | Refer to the discussion of Fig. 3 | $5 \times 10^9$ |

<sup>a</sup> The diffusivity of Aβ monomers is $0.623 \times 10^{-6}$ cm<sup>2</sup>/s, according to ref. [29]. This value is lower than the estimate provided in ref. [42], which is $1.43 \times 10^{-6}$ cm<sup>2</sup>/s, as ref. [29] incorporates the effects of tortuosity. Tortuosity accounts for the decreased diffusion rates in the brain tissue caused by obstacles such as cells and the extracellular matrix.
<sup>b</sup> The values reported in ref. [18] were obtained by curve-fitting experimental data from ref. [31]. The specific parameters are $k_1 = 8 \times 10^{-6}$ h<sup>-1</sup>, $k_2 = 3.4 \times 10^{-2}$ $\mu\text{M}^{-1}$ h<sup>-1</sup>, as indicated in ref. [18].
<sup>c</sup> 0.01 h<sup>-1</sup> [32].
<sup>d</sup> 0.00065 $\mu\text{M}^{-1}$ h<sup>-1</sup> [32].
<sup>e</sup> Used in calculations as a representative value.
<sup>f</sup> The molecular weight of Aβ 42, as reported in ref. [34].
<sup>g</sup> Depending on the specific isoform [35].
<sup>h</sup> According to ref. [34], the human brain produces Aβ 42 monomers at a rate of $0.33 \times 10^{-6}$ $\mu\text{mol s}^{-1}$ . Given the brain's estimated volume of $1.5 \times 10^{15}$ $\mu\text{m}^3$ , the production rate of Aβ 42 per unit volume is approximately $2.2 \times 10^{-22}$ $\mu\text{mol } \mu\text{m}^{-3} \text{ s}^{-1}$ . Within a small cube of side length $L$ (representing the gray-shaded CV in Fig. 1), the total production rate of Aβ 42 peptide is calculated to be $2.75 \times 10^{-17}$ $\mu\text{mol s}^{-1}$ . Aβ monomers are generated at lipid membranes [43]. Assuming that all Aβ 42 monomers are released
<sup>i</sup> Tau is predominantly synthesized in the soma. In healthy neurons, tau is transported to the axon; however, in AD it is misdirected into the somatodendritic compartment [35,44]. The accumulation of tau monomers in the soma results from two key processes: (i) the synthesis of new tau monomers and (ii) the displacement of tau from the axon, which occurs in AD.
Eq. (21) was used to estimate $q_{\text{tau}}$ using the following parameter values: $\frac{V_{\text{NSF},f}}{V_{\text{soma}}} = 0.081$ ,
<sup>j</sup> According to ref. [36], the preclinical phase of AD can last 20 years. Since most neurons begin to die after this phase, the growth period for senile plaques may last for approximately 20 years or even longer.
<sup>k</sup> 10 years.
<sup>l</sup> According to ref. [29] the half-life of Aβ monomers is approximately 1.28 hours.
<sup>m</sup> The half-life of free Aβ aggregates is expected to exceed that of Aβ monomers; however, these aggregates can be broken down by microglia [45]. For modeling purposes, the half-life of free Aβ aggregates is assumed to be ten times that of Aβ monomers.

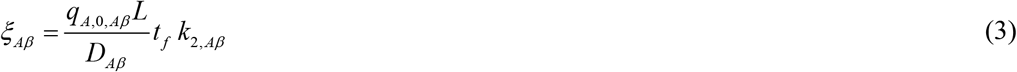

where *q*_*A*,0, *Aβ*_ represents the flux of Aβ monomers across the lower boundary of the shaded CV shown in Fig. 1, *L* is half the distance between senile plaques, *D*_*Aβ*_ denotes the diffusivity of Aβ monomers, *t*_*f*_ is the growth period of senile plaques leading up to neuronal death, and *k*_2, *Aβ*_ is the autocatalytic rate constant for the second step in the F-W model describing Aβ aggregation.

**Fig. 1.**
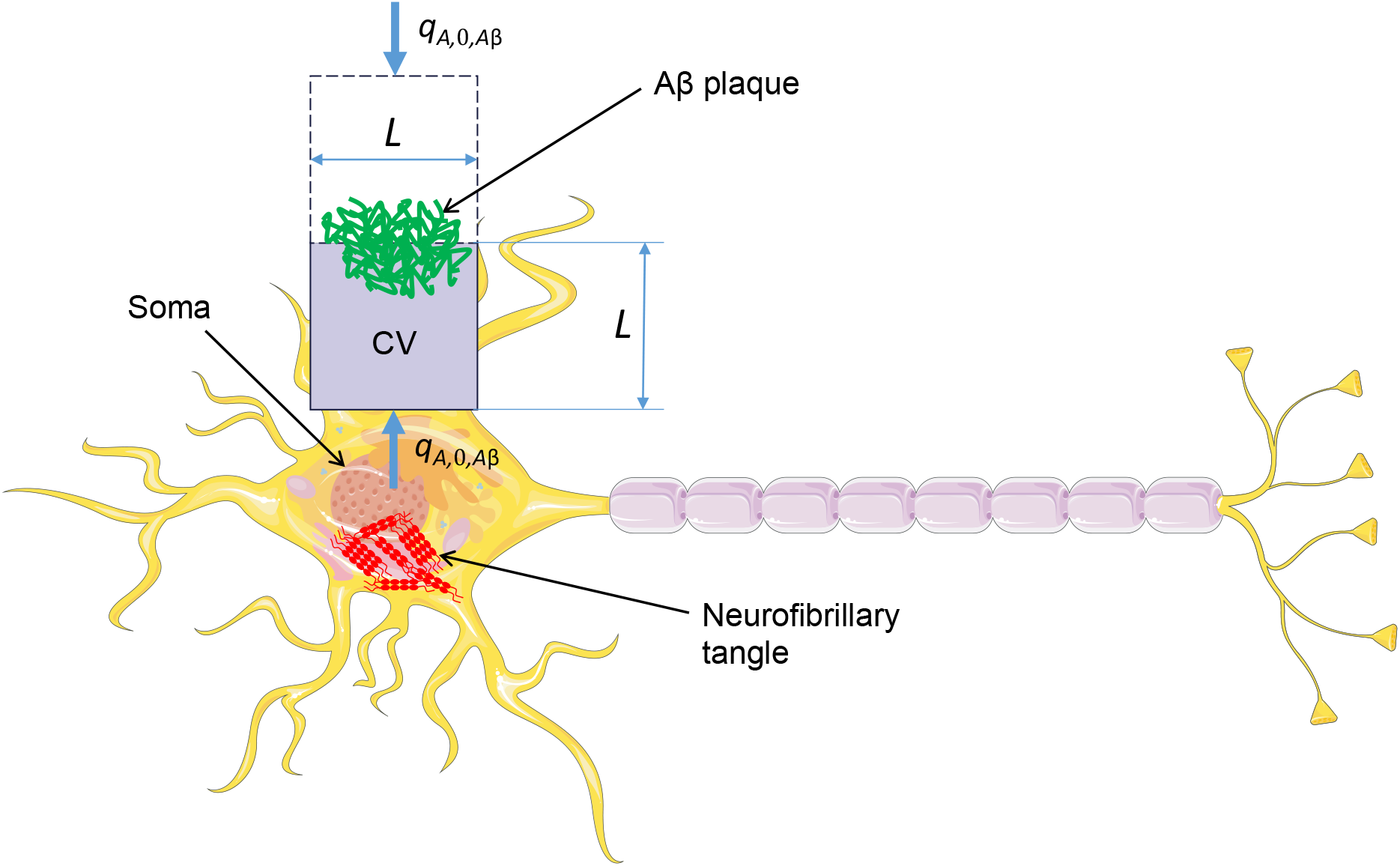
Schematic diagram of a neuron. The figure illustrates an Aβ plaque, a cubic volume in the extracellular space, and neurofibrillary tangle within the soma. Aβ monomers are generated at lipid membranes, with the model positing that these monomers are produced at the lower face of the cubic CV, which has an area of *L*^2^. Tau monomers are synthesized in the soma and also enter the soma due to the displacement of tau from the axon in AD. The formation of an Aβ plaque is modeled to occur symmetrically at the interface between two adjacent, identical CVs, with each CV assumed to contain half of the plaque. The lumped capacitance approximation is employed, assuming that Aβ and tau concentrations are spatially uniform within their respective CVs. Consequently, all Aβ and tau concentrations, including those of monomers and aggregates, are treated as functions of time only.

The lumped capacitance approximation is a useful approach in the transient heat conduction analysis, as it assumes a spatially uniform temperature distribution within the CV, making the temperature solely a function of time [25]. This study focuses on Aβ and tau protein concentrations; therefore, the lumped capacitance approximation assumes spatial uniformity of concentrations, making them dependent only on time.

Ref. [24] showed that the lumped capacitance approximation is valid for the CV when the condition *ξ*_*Aβ*_ << 1 is met, allowing Aβ concentrations to be treated as time-dependent only, with no spatial variation within the CV. The estimation of *ξ*_*Aβ*_ was based on the following parameter values given in Table 2: *D*_*Aβ*_ = 62.3 μm^2^/s, *L*=50 μm, *q*_*A*, 0,*Aβ*_ = 1.1×10^−5^ μM μm s^-1^, *t*_*f*_ = 6.31×10^8^ s, and *k*_2,*Aβ*_ = 9.44 ×10^−6^ μM^-1^ s^-1^. Substituting these values into Eq. (3) yields *ξ*_*Aβ*_ = 0.053. This is enough to consider *ξ*_*Aβ*_ <<1 satisfied, making it possible to neglect any spatial variations in Aβ monomer concentration within the CV. Consequently, time *t* becomes the only independent variable in the model. By employing the principle of conservation for Aβ monomers within the gray-shaded CV shown in Fig. 1, the following equation is obtained:

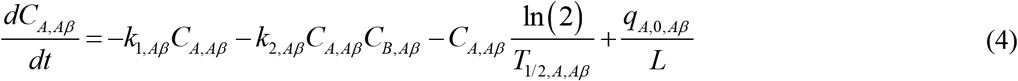

The first term on the right side of Eq. (4) represents the conversion rate of Aβ monomers into free aggregates driven by nucleation, while the second term represents the conversion rate through autocatalytic growth of aggregates. The third term accounts for the degradation of Aβ monomers, and the fourth term represents the generation of Aβ monomers on lipid membranes. Given that *q*_*A*,0, *Aβ*_ denotes the flux of Aβ monomers per unit area at the lower boundary of a CV with a volume *L*^3^, the rate of Aβ monomer production per unit volume in the lumped capacitance model can be expressed as *q*_*A*, 0,*Aβ*_ *L*^2^ / *L*^3^, hence the value of the fourth term in Eq. (4).

By applying the conservation principle to free Aβ aggregates within the CV, the following equation is obtained:

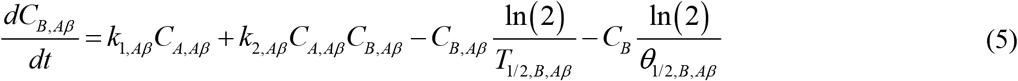

In Eq. (5), the first two terms on the right-hand side mirror the first two terms in Eq. (4) but with opposite signs. The third term indicates the degradation rate of free Aβ aggregates, while the fourth term represents the rate at which these aggregates deposit into senile plaques.

The process by which Aβ plaques form from Aβ aggregates is modeled in a manner akin to colloidal suspension coagulation, as described in ref. [26]. In this approach, free Aβ aggregates (denoted as *B*) are assumed to deposit into Aβ plaques, with the concentration of these deposited aggregates denoted as *C*_*D, Aβ*_:

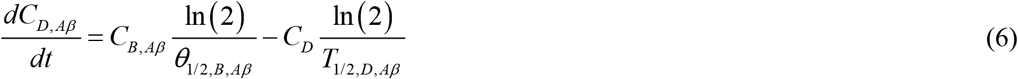

In Eq. (6), the first term on the right-hand side is analogous to the fourth term in Eq. (5) but with the opposite sign. The second term models the possible degradation of Aβ aggregates within the senile plaques.

The initial conditions are as follows:

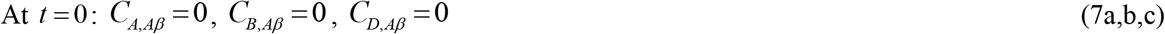

### 2.2. Equations simulating aggregation of tau protein

When tau monomers diffuse rapidly, their concentration, *C*_*A,tau*_, within the soma (Fig. 1) becomes uniform. In this case, time *t* serves as the sole independent variable in the model. Since tau aggregation differs significantly from that of Aβ, as tau aggregates form within neurons, the validity analysis of the lumped capacitance approximation presented in ref. [24] does not apply to tau. This analysis must be revisited for the new context, which is done in Section 2.4.

The F-W model, as defined by Eqs. (1) and (2), is employed once again. In this framework, *A* denotes tau monomers, while *B* denotes free tau aggregates, including dimers, oligomers, and protofibrils. Hereafter, the terms “free tau aggregates” and “oligomers” are used interchangeably, although, in the context of the F-W model, aggregates have a slightly broader meaning than just oligomers. The influx of tau monomers into the soma, denoted as *q*_*tau*_, results from two processes: the synthesis of new tau in the soma and the displacement of tau from the axon, a phenomenon observed in AD. Therefore, the term “supply” is favored over “production” for tau monomers. It is assumed that the supply of tau monomers into the soma occurs at a constant rate, *q*_*tau*_.

Applying the principle of conservation for tau monomers in the soma (Fig. 1) yields:

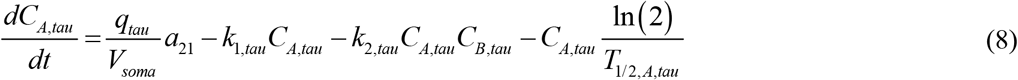

where *C*_*A,tau*_ represents the concentration of tau monomers in the soma, *C*_*B,tau*_ is the concentration of tau oligomers, *k*_1,*tau*_ is the nucleation rate constant for tau oligomers, *k*_2,*tau*_ represents the autocatalytic growth rate constant, and *T*_1/ 2, *A,tau*_ denotes the half-life of tau monomers.

In Eq. (8), the first term on the right-hand side describes the supply of tau monomers, while the second and third terms represent their conversion into tau aggregates through nucleation and autocatalysis, respectively. The fourth term represents the decay of tau monomers because of their finite half-life.

Applying the conservation principle to free tau aggregates results in

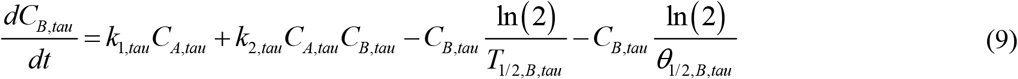

where *θ*_1/ 2, *B, AS*_ denotes the half-deposition time of free tau aggregates into NFTs.

In Eq. (9), the first two terms describe the increase in free tau aggregate concentration resulting from nucleation and autocatalytic growth. The third term accounts for the degradation of free tau aggregates, while the fourth term represents the decrease in free tau aggregates due to their deposition into NFTs. Eqs. (8) and (9) are analogous to Eqs. (3) and (4), respectively.

The formation of NFTs from free tau aggregates within the soma is again modeled similarly to the coagulation process in colloidal suspensions, as described in ref. [26]. Here, free tau aggregates (*B*) are assumed to deposit into NFTs, with the concentration of these deposited aggregates represented by *C*_*D,tau*_. This leads to an expression analogous to Eq. (5):

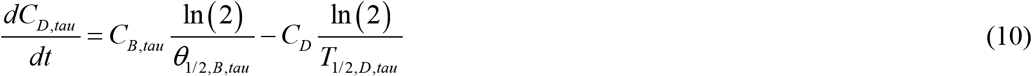

where *T*_1/2, *D,tau*_ represents the half-life of the deposited tau aggregates. In Eq. (10), the first term on the right-hand side mirrors the fourth term in Eq. (9) but with an opposite sign, indicating the deposition of free tau aggregates. The second term models the degradation of tau aggregates within NFTs, assuming they have a finite half-life. This term accounts for possible future therapies, such as the clearance of pathological tau species using antibodies [27,28].

Eqs. (8)–(10) are solved subject to the following initial conditions:

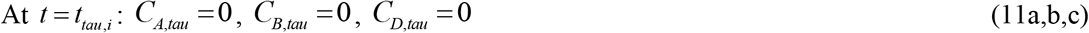

### 2.3. Criterion of combined accumulated toxicity of Aβ and tau oligomers

This paper proposes evaluating the accumulated toxicity resulting from the combined effects of Aβ and tau oligomers using the following expression:

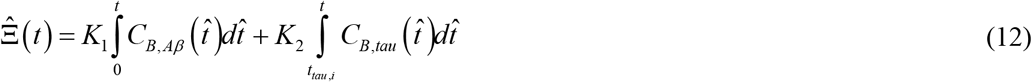

where *t*_*tau, i*_ = 10 years [11]. Since accumulated toxicity can be rescaled arbitrarily, Eq. (12) can be normalized by dividing it by *K*_1_, resulting in

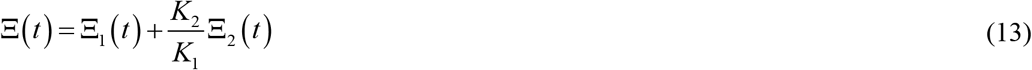

where

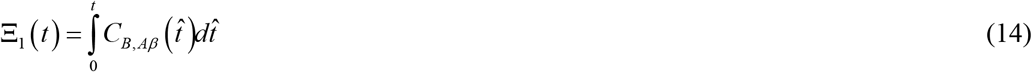

and

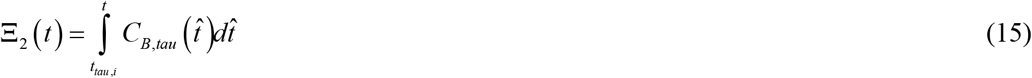

Section S1 of the Supplementary Materials gives analytical solutions for Aβ peptide aggregation for the limiting cases of slow and fast deposition rates of free Aβ aggregates into plaques [48-50]. Similarly, Section S2 gives analytical solutions for tau protein aggregation under the conditions of slow and fast deposition rates of free tau aggregates into NFTs.

### 2.4. Criterion for the validity of the lumped capacitance approximation for tau protein

Tau aggregation differs significantly from Aβ aggregation due to tau’s localization within neurons. As a result, the lumped capacitance approximation analysis presented in ref. [24] for Aβ cannot be directly applied to tau and must be specifically reevaluated for tau aggregation. To conduct this analysis, Eq. (8) requires an additional term to account for the diffusion of tau monomers. Reformulating Eq. (8) within a spherical coordinate system, and assuming a perfectly spherical neuron soma, the conservation of tau monomers can be expressed as:

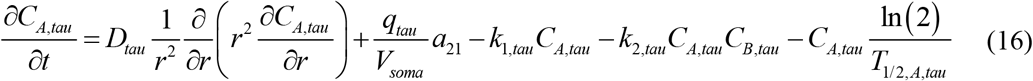

The criterion for assessing the validity of the lumped capacitance approximation is formulated within the context of impaired degradation mechanisms for tau monomers and oligomers, i.e., *T*_1/ 2, *A*_ →∞ and *T*_1/ 2,*B*_ →∞. Applying this approximation and summing Eqs. (9) and (16) under steady-state conditions yields:

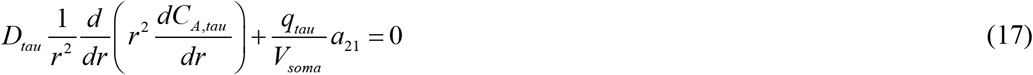

The solution of Eq. (17) is obtained under the following boundary conditions. In the center of the soma, a symmetry condition is utilized:

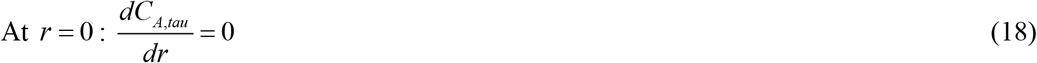

Furthermore, a concentration of tau monomers is postulated at the soma membrane (the wall):

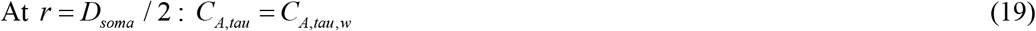

The solution of Eqs. (17)–(19) yields the following concentration profile of tau monomers in the soma:

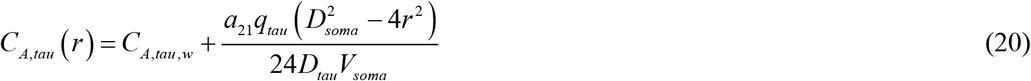

From Eq. (20), it follows that

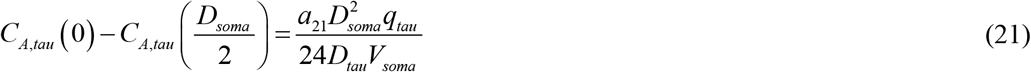

For a slow deposition rate of free tau oligomers into NFTs, *θ*_1/2,*B,tau*_ →∞, the approximate solution for *C*_*A*_ is given by Eq. (S24) of the Supplemental Materials. When considering large time values, Eq. (S24) simplifies to

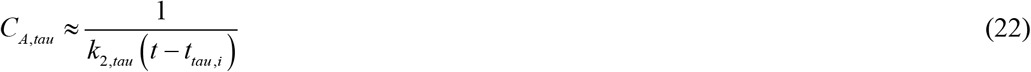

By employing Eqs. (21) and (22), the following parameter is obtained to represent the ratio of the variation of *C*_*A*_ between the center of the soma and the soma membrane relative to the average value of *C*_*A,tau*_ within the soma at time *t*_*f*_ :

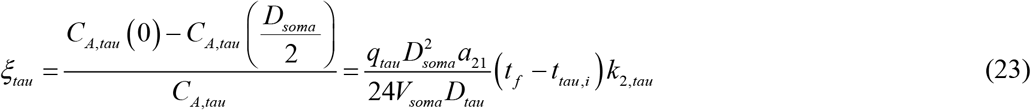

The role of parameter *ξ*_*tau*_ is similar to that of the Biot number in convection-conduction problems. The lumped capacitance approximation is valid as long as *ξ*_*tau*_ << 1. Importantly, the error in the lumped capacitance approximation increases as the process duration *t* − *t*_*tau, i*_ lengthens.

The estimation of *ξ* was based on the following parameter values: *D*_*tau*_ = 3 μm^2^/s, *q*_*tau*_ = 3.63×10^−23^ mol s^-1^, *t*_*tau. i*_ = 3.15×10^8^ s, *k*_*2, tau*_ = 1.81×10^−7^ μM^-1^ s^-1^, *D*_*soma*_ = 20 μm, *V*_*soma*_ = 4.19 ×10^3^ μm^3^, and 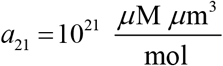. Substituting these values into Eq. (23) results in *ξ* = 0.0028. This is enough for the criterion *ξ*_*tau*_ << 1 to be satisfied, indicating that variations in tau concentration across the soma can be neglected.

A model for the growth of senile plaques is presented in Section S3 of the Supplementary Materials.

### 2.5. Method for calculating the volume occupied by NFTs

The increase in NFT volume (Fig. 1) is determined by calculating the total number of tau monomers incorporated into NFTs within the soma, denoted as *N*_*tau*_, over a specified time *t*. Applying the approach adapted from ref. [51] yields:

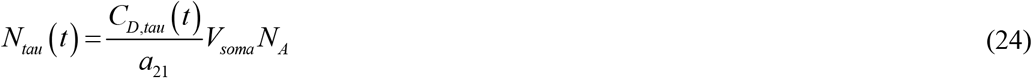

where *N*_*A*_ is Avogadro’s number.

Alternatively, *N*_*tau*_ (*t*) may be calculated using the following equation, as described in ref. [51]:

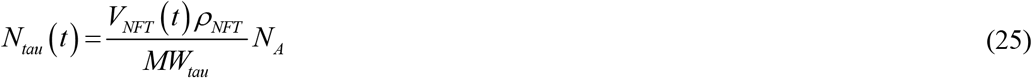

where *MW*_*tau*_ denotes the molecular weight of a tau monomer, *V*_*tau*_ (*t*) represents the NFT volume in the soma at time *t*, and *ρ*_*NFT*_ is the NFT density.

Equating the right-hand sides of Eqs. (24) and (25) and solving for the NFT volume gives:

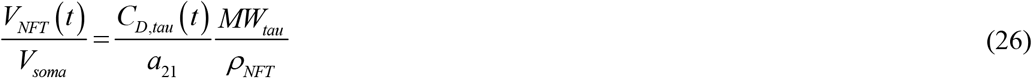

Substituting the expression for *C*_*D, tau*_ from Eq. (S34) into Eq. (26) results in

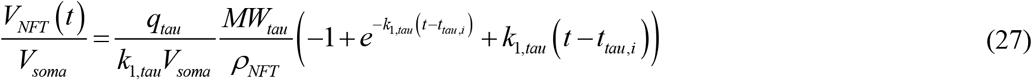

Using Eq. (27) at *t* = *t*_*f*_ and solving it for *q*_*tau*_ yields:

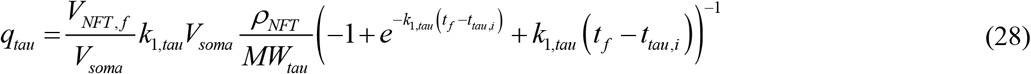

Eq. (28) is crucial for estimating model parameters because it enables the calculation of *q*_*tau*_ when the values of 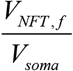, *t*_*tau, i*_, *t*_*f*_, and *k*_1, *tau*_ are known. Using 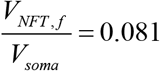, *t*_*tau, i*_ = 3.15×10^8^ s, *t*_*f*_ = 6.31×10^8^ s, and *k*_1,*tau*_ = 2.78×10^−6^ s^-1^ in Eq. (28), the value of *q*_*tau*_ is estimated to be 1.57×10^−28^ mol s^−1^, as listed in Table 2.

## 3. Results

Details of the numerical solution are given in Section S4 of the Supplementary Materials. Parameter values given in Table 2 are used, except where indicated otherwise in the figure or its accompanying caption. The simulation timescale is based on the estimated 20-year growth period of senile plaques [36], which aligns with the duration of the preclinical phase of AD preceding the onset of widespread neuronal death.

Once the aggregation of Aβ and tau begins, the concentrations of free aggregates quickly reach their equilibrium values (Fig. 2). The observed higher concentration of tau aggregates compared to Aβ aggregates is attributed to the differences in the equilibrium molar concentrations of free Aβ and tau aggregates (Fig. 2). Note that tau aggregation begins at *t* = *t*_*tau, i*_ (10 years in this case). In the scenario involving a dysfunctional protein degradation machinery, characterized by infinitely long half-lives for monomers and free aggregates, the equilibrium concentrations of free tau aggregates are unaffected by the nucleation rate constants for these aggregates, denoted as *k*_1,*Aβ*_ and *k*_1,*tau*_ (Fig. 2b). A similar trend is observed for free Aβ aggregates when Fig. 2b is replotted with a larger *y*-axis scale (see Fig. S1b). In contrast, when considering physiologically relevant half-lives for monomers and free aggregates, the equilibrium concentration of tau becomes dependent on *k*_1,*tau*_. A higher value of *k*_1,*tau*_ leads to a greater concentration of free tau aggregates due to the accelerated conversion of tau monomers into aggregates (Fig. 2a). A similar pattern is observed for free Aβ aggregates when Fig. 2a is replotted with a larger *y*-axis scale: higher values of *k*_1,*Aβ*_ correspond to increased concentrations of free Aβ aggregates (see Fig. S1a). Furthermore, the concentrations of Aβ and tau aggregates are several times larger in the scenario of dysfunctional protein degradation machinery compared to the scenario with physiologically relevant half-lives for Aβ and tau monomers and free aggregates (see Figs. 2a and 2b as well as Figs. S1a and S1b).

**Fig. 2.**
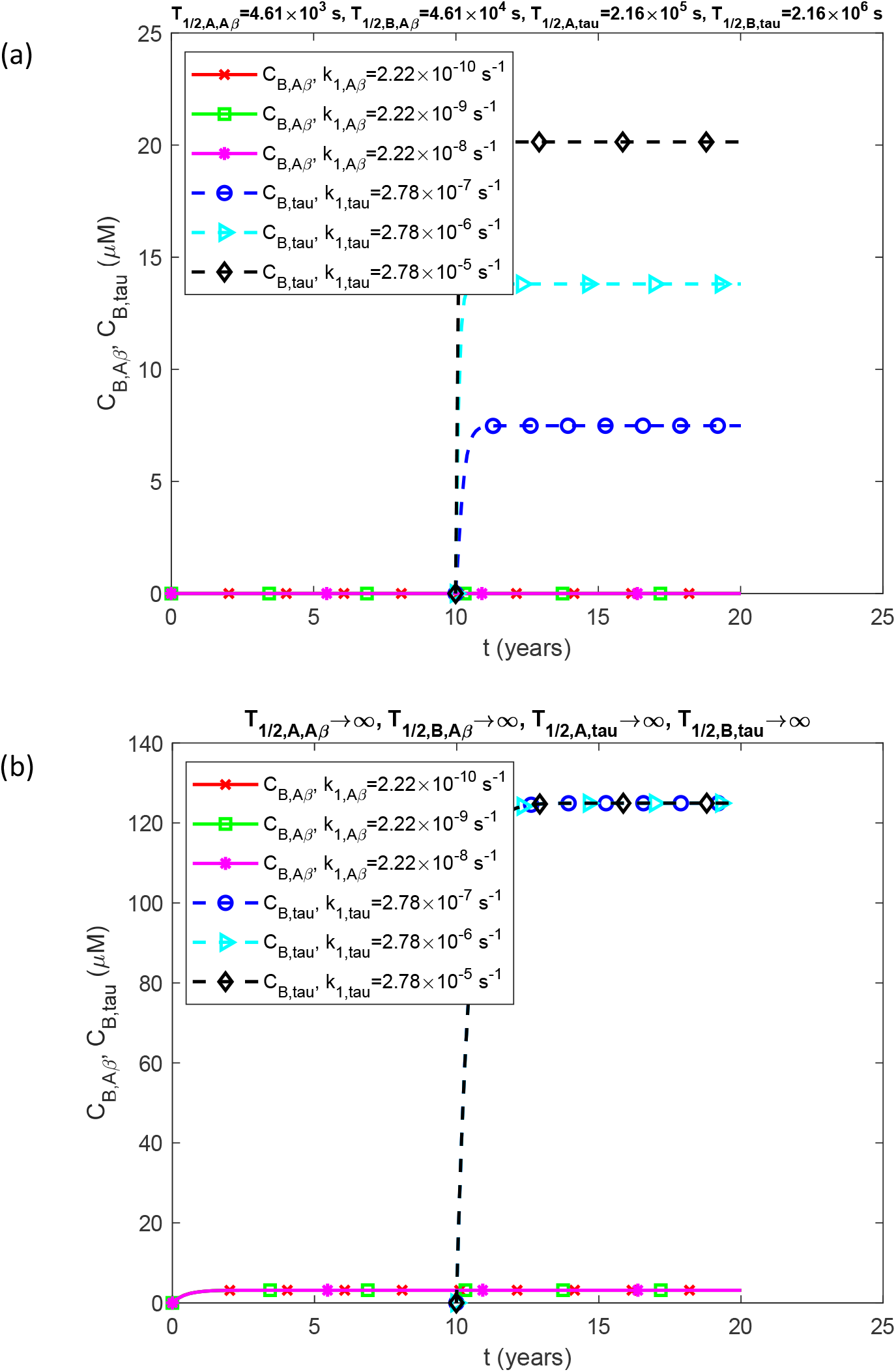
Molar concentrations of free Aβ aggregates (oligomers), *C*_*B, Aβ*_, and free tau aggregates (oligomers), *C*_*B,tau*_, as functions of time (in years). *C*_*B, Aβ*_ and *C*_*B,tau*_ are solutions of differential equations (4)-(6) and (8)-(10) and are therefore influenced by model parameters, such as the half-lives of Aβ and tau monomers and aggregates (see Table 2). (a) This scenario represents physiologically relevant half-lives of Aβ and tau monomers and aggregates. (b) This scenario illustrates dysfunctional protein degradation machinery corresponding to infinitely long half-lives of Aβ and tau monomers and aggregates. Results are presented for three different values of nucleation rate constants for free Aβ and tau aggregates, *k*_1,*Aβ*_ and *k*_1,*tau*_, respectively. Parameters: *T*_1/2,*D,Aβ*_ →∞, *T*_1/2,*D,tau*_ →∞, *θ* _1/2,*B,Aβ*_ =10^7^ s, and *θ* _1/2,*D,tau*_ =10 s.

The influence of autocatalytic growth rate constants for free Aβ and tau aggregates, *k*_2, *Aβ*_ and *k*_2,*tau*_, is illustrated in Fig. S2 in the Supplemental Materials, and it is similar to the effect of nucleation rate constants. In the scenario with physiologically relevant half-lives for monomers and free aggregates, the concentration of tau aggregates rises with increasing values of *k*_2,*tau*_ since higher values of *k*_2,*tau*_ correspond to a more rapid conversion of tau monomers into free aggregates (Fig. S2a). Conversely, when considering infinite half-lives for monomers and aggregates, the concentration of tau aggregates remains unaffected by changes in *k*_2,*tau*_ (Fig. S2b).

Analysis of Figs. 2 and S1 suggests that at high concentrations of free tau aggregates (as shown in Figs. 2b and S1b) the production of free tau aggregates is not limited by kinetic processes. This is due to the autocatalytic nature of tau monomer conversion into free aggregates. Thus, in these high-aggregate concentration scenarios, tau levels appear to be constrained by the supply rate of tau monomers to the soma. This hypothesis is further corroborated by data in Fig. S3, which demonstrates that an increased supply rate of tau monomers into the soma leads to a rise in the concentration of tau aggregates, regardless of whether the half-lives of tau monomers and aggregates are finite or infinite (Figs. S3a and S3b).

The combined accumulated toxicity of Aβ and tau oligomers increases linearly over time (Fig. 3) due to the relatively constant concentrations of free Aβ and tau aggregates (Fig. 2). Since accumulated toxicity is calculated by integrating oligomer concentrations over time (see Eqs. (14) and (15)), it has a direct linear dependence on time. When considering finite half-lives for monomers and free aggregates, accumulated toxicity is affected by the values of *k*_1,*Aβ*_ and *k*_1,*tau*_ (Fig. 3a). However, with infinite half-lives of monomers and free aggregates, accumulated toxicity becomes independent of both *k*_1,*Aβ*_ and *k*_1,*tau*_, in agreement with the findings in Fig. 2. A similar trend is evident in Fig. S4, which presents accumulated toxicity for varying values of *k*_2, *Aβ*_ and *k*_2,*tau*_, consistent with results shown in Fig. S2. For both finite and infinite half-life scenarios, accumulated toxicity rises with increasing values of *q*_*A*,0, *Aβ*_ and *q*_*tau*_ (Fig. S5), consistent with the trends shown in Fig. S3.

**Fig. 3.**
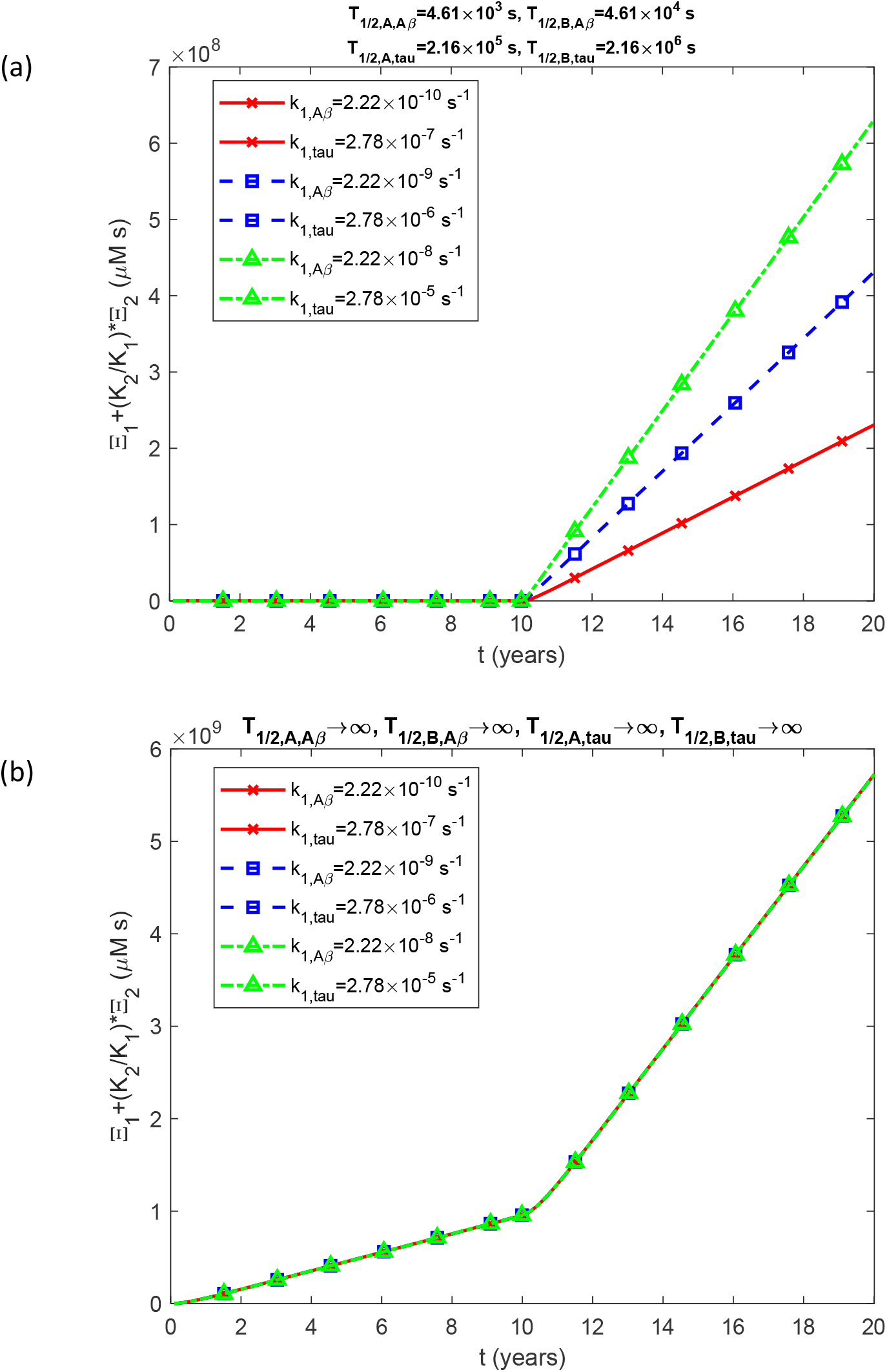
Combined accumulated toxicity of Aβ and tau oligomers, 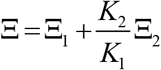, as a function of time (in years). (a) This scenario illustrates physiologically relevant half-lives of Aβ and tau monomers and aggregates. (b) This scenario depicts dysfunctional protein degradation machinery, characterized by infinitely long half-lives of Aβ and tau monomers and aggregates. Results are presented for three different values of nucleation rate constants for free Aβ and tau aggregates, *k*_1,*Aβ*_ and *k*_1,*tau*_, respectively. Parameters: *T*_1/2,*D,Aβ*_ →∞, *T*_1/2,*D,tau*_ →∞, *θ* _1/2,*B,Aβ*_ =10^7^ s, and *θ*_1/2,*B,tau*_ =10^7^ s.

It is widely accepted that AD is linked to dysfunctional degradation mechanisms for Aβ and tau proteins [52-54]. In the developed model, such dysfunction is represented by assigning infinite half-lives to Aβ and tau monomers and free aggregates. Based on Figs. 3b and S3b, with physiologically realistic rates of Aβ and tau monomer production and supply, the critical threshold of combined accumulated toxicity from Aβ and tau oligomers (the level at which neuron death occurs) is estimated to be 5 ×10^9^ µM·s.

In plotting Figs. 4, S6, and S7, parameters *θ*_1/ 2,*B, Aβ*_ and *θ*_1/2,*B,tau*_ were set to equal values and increased simultaneously. The combined accumulated toxicity, plotted against the half-deposition times of free Aβ aggregates into senile plaques (*θ*_1/2,*B, Aβ*_) and tau aggregates into NFTs (*θ*_1/2,*B,tau*_), follows an S-shaped curve. The S-shaped trend, shown in Fig. 4, suggests that senile plaques and NFT formation may reduce neurotoxicity caused by Aβ and tau oligomers. Rapid deposition of Aβ and tau oligomers into senile plaques and NFTs corresponds to shorter half-deposition times of free Aβ and tau aggregates, allowing these oligomers to be quickly sequestered into insoluble aggregates and resulting in minimal oligomer-induced toxicity (Fig. 4). The hypothesis that the formation of plaques and tangles may reduce oligomer-induced toxicity is further supported by comparing Fig. 4 with Figs. 5a and 6a, where smaller values of *θ*_1/2,*B, Aβ*_ and *θ*_1/2,*B,tau*_ are associated with larger plaques and NFTs. Furthermore, accumulated toxicity reaches its highest levels at longer half-deposition times (Fig. 4), while Figs. 5a and 6a show that larger values of *θ*_1/2,*B, Aβ*_ and *θ*_1/2,*B,tau*_ correspond to smaller plaques and NFTs.

**Fig. 4.**
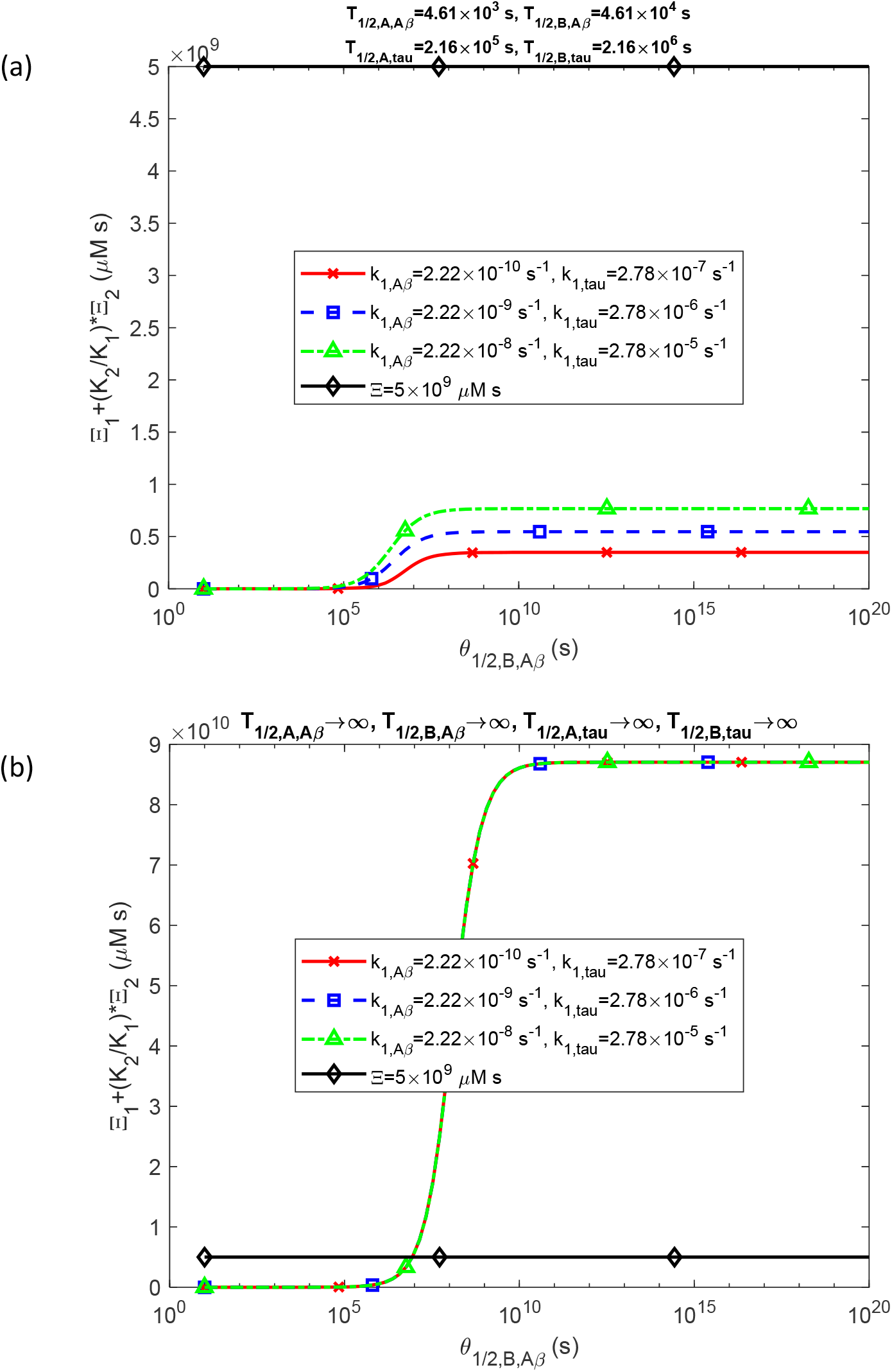
Combined accumulated toxicity of Aβ and tau oligomers, 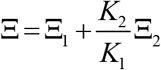, as a function of the half-deposition times, *θ*_1/2,*B,Aβ*_ and *θ*_1/2,*B,tau*_ (both set to equal values for this computation). (a) This scenario represents physiologically relevant half-lives of Aβ and tau monomers and aggregates. (b) This scenario depicts dysfunctional protein degradation machinery, characterized by infinitely long half-lives of Aβ and tau monomers and aggregates. Results are presented at *t* = *t*_*f*_ = 6.31 × 10^8^ s for three different values of nucleation rate constants for free Aβ and tau aggregates, *k*_1,*Aβ*_ and *k*_1,*tau*_, respectively. Parameters: *T*_1/2,*D,Aβ*_ →∞, *T*_1/2,*D,tau*_ →∞.

**Fig. 5.**
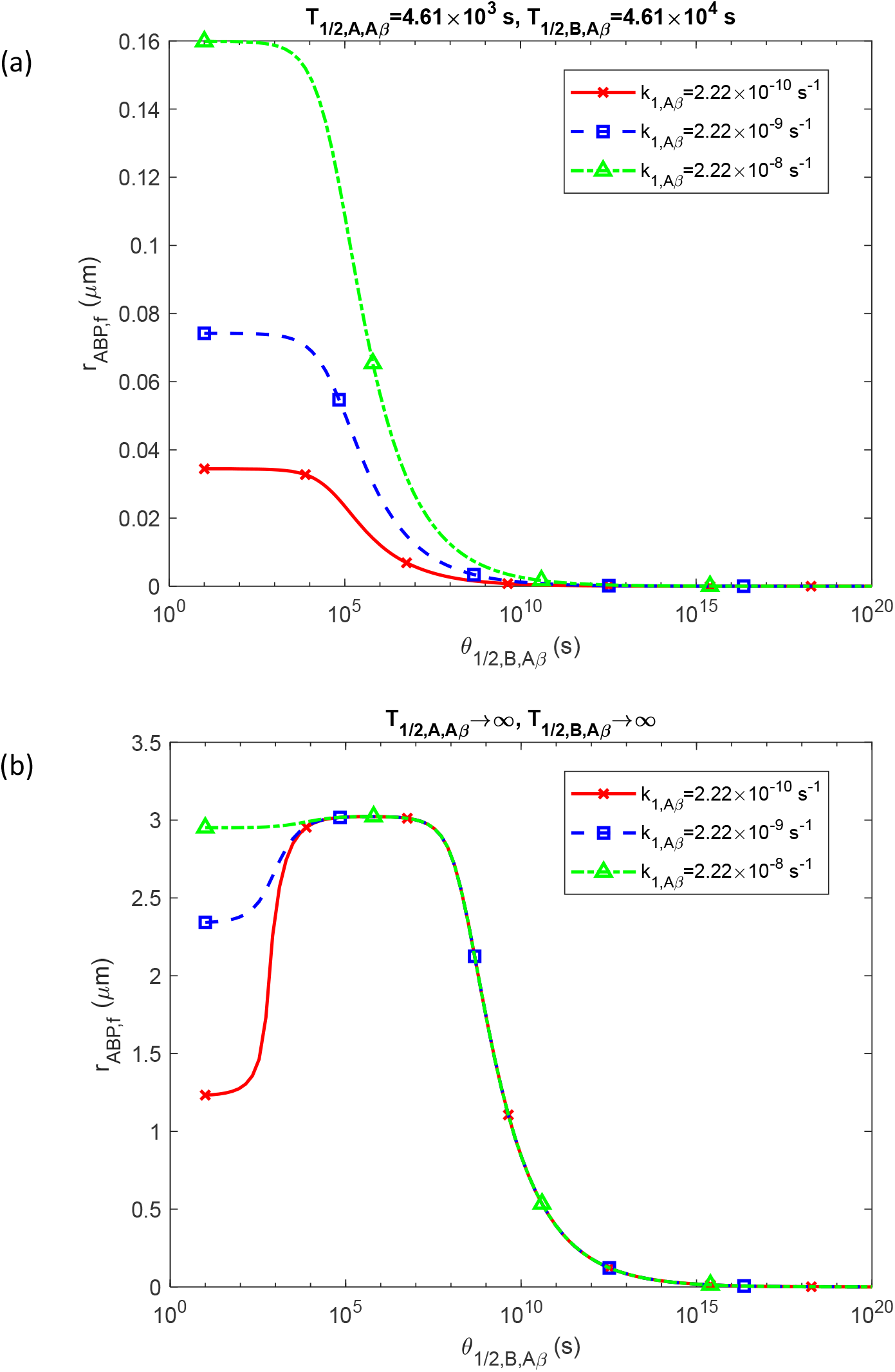
Radius of the growing Aβ plaque, *r*_*ABP, f*_, at *t* = *t*_*f*_ shown as a function of the half-deposition time *θ*_1/2,*B,Aβ*_. (a) Scenario with physiologically relevant half-lives for Aβ monomers and aggregates. (b) Scenario illustrating dysfunctional protein degradation machinery, with infinitely long half-lives for Aβ monomers and aggregates. Results are presented for three different nucleation rate constants for free Aβ aggregates, *k*_1,*Aβ*_. Parameters: *t* _*f*_ = 6.31 × 10^8^ s, *T*_1/2,*D,Aβ*_ →∞.

Notably, when the protein degradation machinery is functional—reflecting physiologically relevant half-lives for Aβ and tau monomers and free aggregates—the accumulated toxicity remains below the critical threshold (Fig. 4a). However, with infinite half-lives for monomers and aggregates, accumulated toxicity can exceed the critical level by nearly twentyfold as *θ*_1/ 2,*B, Aβ*_ and *θ*_1/2,*B,tau*_ increase (Fig. 4b). The nucleation rates *k*_1,*Aβ*_ and *k*_1,*tau*_ impact accumulated toxicity only when half-lives of monomers and aggregates are finite; in this case, increasing *k*_1,*Aβ*_ and *k*_1,*tau*_ results in higher accumulated toxicity (Fig. 4a).

For infinitely long half-lives of monomers and aggregates, the maximum size of an Aβ plaque is observed when *θ*_1/2,*B, Aβ*_ ranges between approximately 10^5^ and 10^7^ s (Fig. 5b). This is attributed to the role of autocatalysis in converting Aβ monomers into free aggregates. For low *θ*_1/2,*B, Aβ*_ values, free Aβ aggregates rapidly deposit into plaques, resulting in a low concentration of free Aβ aggregates and consequently a reduced rate of autocatalytic conversion of Aβ monomers into aggregates. This leads to fewer Aβ aggregates available for deposition into Aβ plaques. Conversely, at high *θ*_1/ 2,*B, Aβ*_ values, free aggregates remain in suspension rather than depositing into plaques, producing the largest plaques at intermediate *θ*_1/ 2,*B, Aβ*_ values.

Fig. 5b demonstrates the effect of the nucleation rate constant, *k*_1,*Aβ*_, on the plaque size. At high *k*_1,*Aβ*_, nucleation proceeds quickly, so the absence of autocatalysis at low *θ*_1/2,*B, Aβ*_ (because of the immediate deposition of aggregates into plaques) does not significantly reduce the plaque size (a green line marked by triangles in Fig. 5b), as all synthesized Aβ monomers can still convert into free aggregates via nucleation. However, at low *k*_1,*Aβ*_, plaque size reduction becomes pronounced as *θ*_1/2,*B, Aβ*_ decreases (a red line marked by crosses in Fig. 5b). In Fig. 6b, the lack of decrease in NFT volume at low *θ*_1/2,*B,tau*_ can be explained by the sufficiently rapid conversion of tau monomers into aggregates through nucleation.

**Fig. 6.**
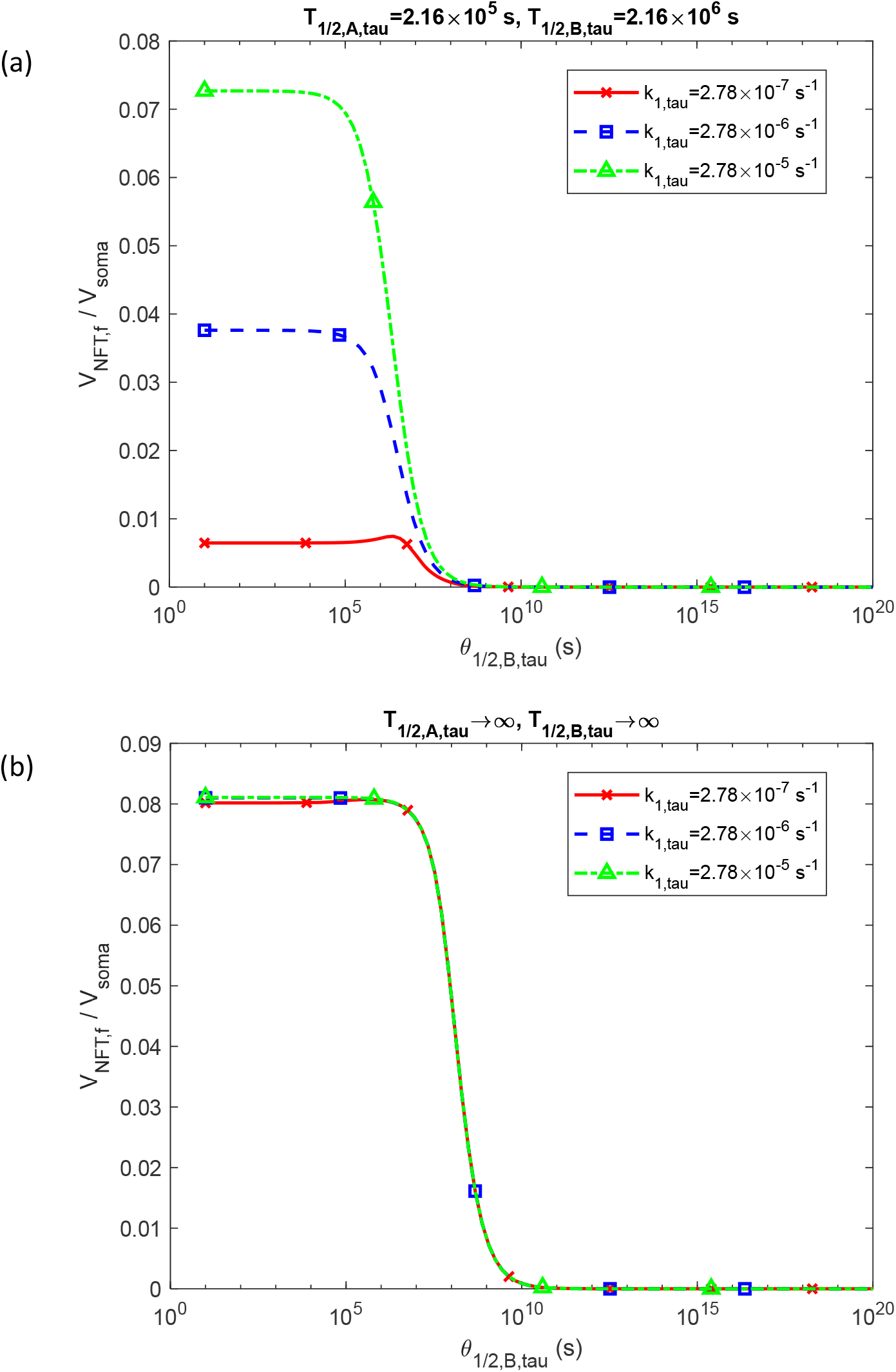
Fraction of the soma volume occupied by NFTs, 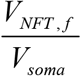, at *t* = *t* _*f*_ shown as a function of the half-deposition time *θ*_1/2,*B,Aβ*_. (a) Scenario representing physiologically relevant half-lives for tau monomers and aggregates. (b) Scenario illustrating dysfunctional protein degradation machinery, with infinitely long half-lives for tau monomers and aggregates. Results are displayed for three different nucleation rate constants for free tau aggregates, *k*_1,*Aβ*_. Parameters: *t* _*f*_ = 6.31 × 10^8^ s, *T*_1/2,*D,tau*_ →∞.

The effect of *k*_2, *Aβ*_ and *k*_2,*tau*_ on the change of accumulated toxicity with respect to *θ*_1/2,*B, Aβ*_ and *θ*_1/ 2,*B,tau*_ is similar to that of *k*_1,*Aβ*_ and *k*_1,*tau*_ (Fig. S6). For finite half-lives of Aβ and tau monomers and free aggregates, an increase in *k*_2, *Aβ*_ raises accumulated toxicity due to the higher autocatalytic conversion rates of Aβ and tau monomers into aggregates (Fig. S6a).

Increasing the production and supply rates of Aβ and tau monomers, *q*_*A*,0, *Aβ*_ and *q*_*tau*_, respectively, raises accumulated toxicity. At the highest values of *q*_*A*,0, *Aβ*_ and *q*_*tau*_, accumulated toxicity may exceed the critical threshold of 5 ×10^9^ µM·s, even with finite half-lives of Aβ and tau monomers and aggregates (Fig. S7a). When monomer and aggregate half-lives are infinite, this critical threshold is exceeded at any of the three values of *q*_*A*,0, *Aβ*_ and *q*_*tau*_ used in Fig. S7b, once *θ*_1/2,*B, Aβ*_ and *θ*_1/2,*B,tau*_ reach sufficiently high levels.

An increase in the autocatalytic constant *k*_2, *Aβ*_ results in a larger Aβ plaque size at lower *θ*_1/2,*B, Aβ*_ values (a green line marked by triangles in Fig. S8b), as high *k*_2, *Aβ*_ enables autocatalysis to contribute significantly even at lower concentrations of free Aβ aggregates. Notably, in Fig. S9a, the peak NFT volume rises with increasing autocatalytic rate constant *k*_2,*tau*_ (a green line marked by triangles in Fig. S9a).

Figs. S10 and S11 demonstrate that the production rates of Aβ and tau monomers, *q*_*A*,0, *Aβ*_ and *q*_*tau*_, are key factors influencing Aβ plaque size and NFT volume. Increases in *q*_*A*,0, *Aβ*_ and *q*_*tau*_ significantly increase the maximum values of *r*_*ABP, f*_ and 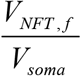.

In computing Fig. 7, *q*_*A*,0, *Aβ*_ and *q*_*tau*_ were increased simultaneously in proportion to their base values given in Table 2: *q*_*A*,0, *Aβ*_ = *aq*_*A*,0, *Aβ, b*_ and *q*_*tau*_ = *aq*_*tau,b*_. Accumulated toxicity rises linearly with the increase of parameter *a*. Since Fig. 7 is plotted in log-log axes, a power-law relationship exists between accumulated toxicity and the production and supply rates of Aβ and tau monomers. Increasing *θ*_1/2,*B, Aβ*_ and *θ*_1/2,*B,tau*_ thus elevates accumulated toxicity.

**Fig. 7.**
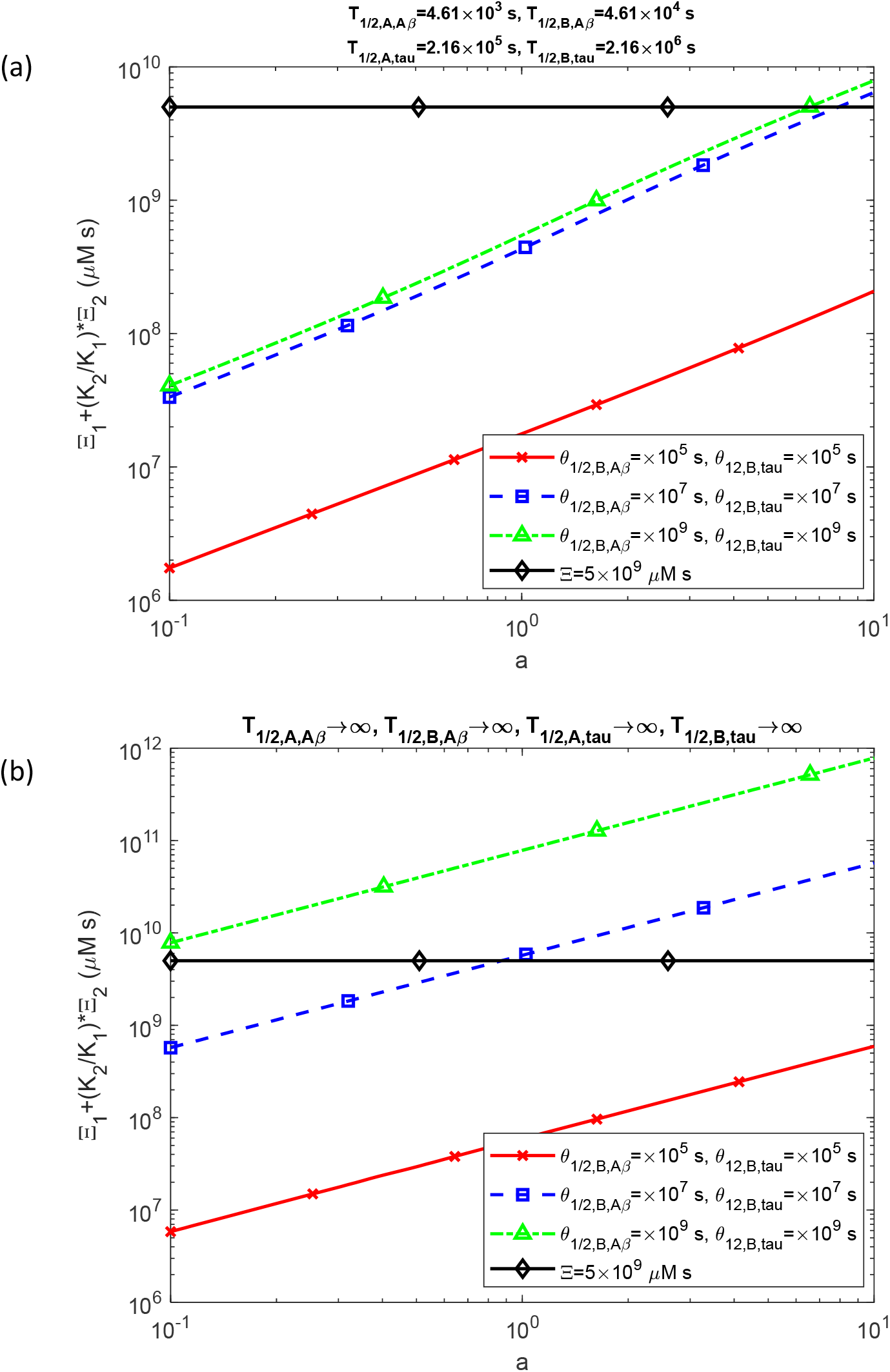
Combined accumulated toxicity of Aβ and tau oligomers, 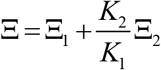, as a function of parameter *a*, which determines the magnitude of production rates for Aβ and tau monomers, *q*_*A*,0, *Aβ*_ and *q*_*tau*_, respectively. In the computation of Fig. 7, *q*_*A*,0, *Aβ*_ and *q*_*tau*_ were calculated as *q*_*A*,0, *Aβ*_ = *aq*_*A*,0, *Aβ, b*_ and *q*_*tau*_ = *aq*_*tau,b*_, where *q*_*A*,0, *Aβ, b*_ and *q*_*tau,b*_ are the values given in Table 2, *q*_*A*,0, *Aβ, b*_ = 1.1×10^−5^ μM μm s^-1^ and *q*_*tau,b*_ = 3.63×10^−23^ mol s^-1^. (a) This scenario depicts physiologically relevant half-lives of Aβ and tau monomers and aggregates. (b) This scenario illustrates dysfunctional protein degradation machinery, characterized by infinitely long half-lives of Aβ and tau monomers and aggregates. Results are presented at *t* = *t* _*f*_ = 6.31 × 10^8^ s for three different values of half-deposition times, *θ*_1/2,*B,Aβ*_ and *θ*_1/2,*B,tau*_. Parameters: *T*_1/2,*D,Aβ*_ →∞, *T*_1/2,*D,tau*_ →∞.

## 4. Conclusions, limitations of the model and future directions

This paper proposed, for the first time, a cellular-level criterion for AD progression, based on the evaluation of accumulated toxicity induced by the combined effects of Aβ and tau oligomers over time. Computational findings indicate that once senile plaques and NFTs begin to form, the concentrations of Aβ and tau oligomers quickly reach equilibrium values and remain constant for the rest of the process. As accumulated toxicity is defined by the integral of oligomer concentration over time, it increases linearly with time. A critical toxicity threshold is estimated, above which neuronal death occurs. Plotting accumulated toxicity versus the half-deposition times of Aβ and tau oligomers into senile plaques and NFTs produces S-shaped curves. Additionally, a power-law relationship is observed between accumulated toxicity and the rates of Aβ and tau monomer production and supply.

The obtained results suggest that senile plaques and NFTs may reduce the toxicity caused by Aβ and tau oligomers by sequestering them into stable, insoluble aggregates.

The half-deposition times of free Aβ and tau aggregates into senile plaques and NFTs are shown to significantly influence the sizes of Aβ and tau inclusions. The largest inclusions occur at intermediate half-deposition times: very short half-deposition times limit the autocatalytic conversion of monomers into free aggregates due to low concentrations of free aggregates, while very long half-deposition times prevent the deposition of free aggregates into these inclusions.

The obtained results suggest that the process of formation of senile plaques and NFTs may be more toxic than the plaques and tangles themselves. The toxicity arises as plaque and NFT formation produces oligomeric intermediates of Aβ and tau, which may be more harmful than the inclusions that eventually form, as noted in ref. [55]. The findings indicate that rapid deposition of Aβ and tau oligomers into senile plaques and NFTs can help reduce overall toxicity by clearing these harmful oligomers from circulation.

These findings align with ref. [15], which observed in experiments that neurons containing tau tangles demonstrated a significantly higher survival rate compared to those without tangles. They are also consistent with data suggesting that NFTs are relatively inert and that soluble oligomeric forms of tau are more toxic to synapses and neurons than NFTs [14]. Ref. [56] also proposed that the formation of Aβ inclusions may act as a neuroprotective mechanism, although one that eventually fails.

The developed model has limitations, as it does not account for how the interactions between the toxic species of Aβ and tau affect accumulated toxicity. It assumes that Aβ and tau aggregates form independently. Given the established cross-talk mechanisms between these processes, such as Aβ-induced tau hyperphosphorylation [57], future research should aim to develop models that capture the interdependent toxicities arising from Aβ and tau oligomers, building on the approaches outlined in refs. [58,59]. Future model development should include time-dependent changes in key parameters, such as protein production rates. Additionally, future research should aim to extend the developed criterion to account for the different toxicities of various oligomer species.

## Abbreviations

Aβ: amyloid beta
AD: Alzheimer’s disease
CV: control volume
F-W: Finke−Watzky
NFT: neurofibrillary tangle

## Data accessibility

This article has no additional data.

## Authors’ contributions

AVK is the sole author of this paper.

## Supplemental Materials

### S1. Analytical solutions for Aβ peptide aggregation in limiting cases

#### S1.1. Analytical solutions for the limiting case of slow Aβ aggregate deposition into senile plaques

In the scenario where the deposition rate of Aβ aggregates into senile plaques is slow and the half-lives of monomers and free aggregates are assumed to be infinitely long, Eqs. (4)–(6) simplify to the following forms:

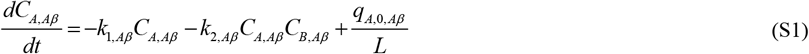

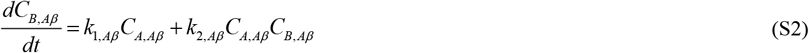

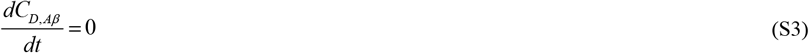

Eqs. (S1) and (S2) were analytically solved with the initial conditions (7a,b), as described in refs. [48,49]. While the exact solution is quite complex, a more streamlined approximate solution can be derived.

Summing Eqs. (S1) and (S2) gives,

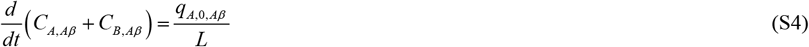

The numerical solution of Eqs. (S1) and (S2) with initial conditions (7a,b) reveals that *C*_*A,Aβ*_ → 0 as *t* →∞. Leveraging this behavior, an approximate solution for *C*_*B, Aβ*_, applicable for large *t*, was obtained in ref. [50] and is expressed as:

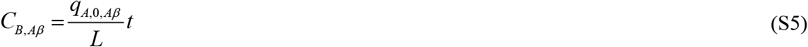

By substituting this expression into Eq. (S2), the following result is obtained:

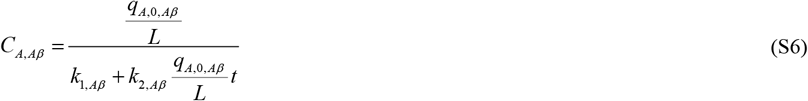

Furthermore, from Eqs. (S3) and (7c), the following is derived:

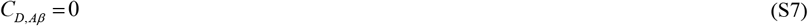

Utilizing Eqs. (14) and (S5) gives

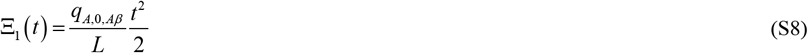

Note that Eq. (S8) is derived for *θ*_1/2,*B,Aβ*_ →∞. This differs from the conditions used to compute Figs. 3, S3, and S4, which are computed for *θ*_1/2,*B,Aβ*_ →10^7^ s (see Table 2).

#### S1.2. Analytical solutions for the scenario of a fast deposition rate of Aβ aggregates into senile plaques, *θ*_1/2,*B,Aβ*_ → 0, as well as *T*_1/2, *A, Aβ*_ →∞, *T*_1/2,*B, Aβ*_ →∞, **and** *T*_1/2,*D, Aβ*_ →∞

The case of *θ*_1/2,*B,Aβ*_ → 0 corresponds to the rapid deposition of free Aβ aggregates into plaques. In this scenario the system of Eqs. (4)–(6) simplifies as follows:

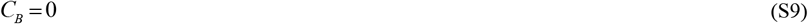

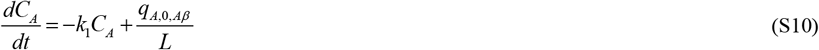

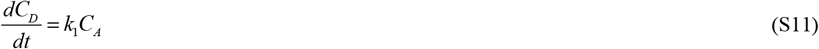

In this scenario, Eqs. (14) and (S9) lead to

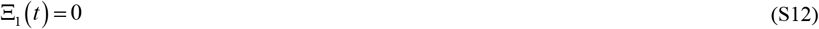

Adding Eqs. (S10) and (S11) yields

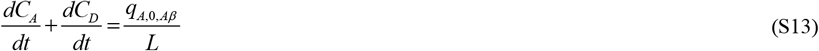

By integrating Eq. (S13) with respect to time and applying initial conditions (7a,c), the following equation is obtained:

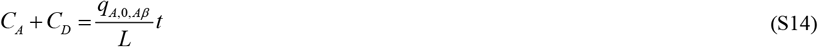

The increase in *C*_*A*_ + *C*_*D*_ over time is explained by the continuous influx of newly synthesized Aβ monomers at the lower boundary of the CV, as illustrated in Fig. 1. Substituting *C*_*A*_ from Eq. (S14) into Eq. (S11) yields the following equation:

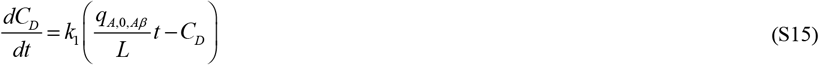

The exact solution of Eq. (S15), subject to the initial condition specified in Eq. (7c), is as follows:

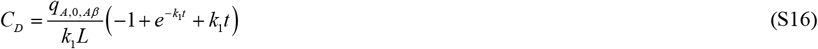

As *t* → 0, Eq. (S16) simplifies to

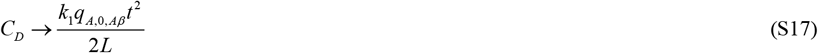

The quantity *C*_*A*_ can then be calculated using Eqs. (S14) and (S16) as follows:

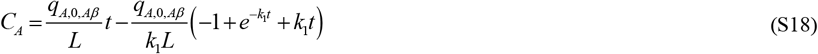

### S2. Analytical solutions for tau protein aggregation in limiting cases

#### S2.1. Analytical solution for the limiting case of slow deposition rate of free tau oligomers into NFTs, *θ*_1/2,*B,tau*_ →∞ (and also *T*_1/ 2, *A,tau*_ →∞, *T*_1/ 2, *B,tau*_ →∞, and *T*1/2, *D,tau* →∞)

In the situation where the deposition rate of free tau aggregates into NFTs is slow, and the half-lives of tau monomers, free aggregates, and deposited aggregates are assumed to be infinitely long, Eqs. (8)–(10) can be simplified to the following forms:

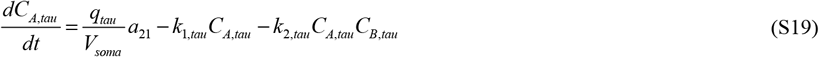

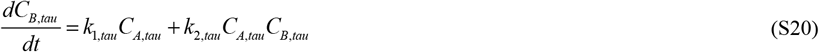

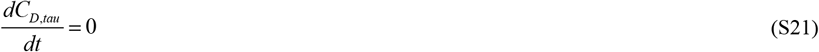

Summing Eqs. (S19) and (S20) gives

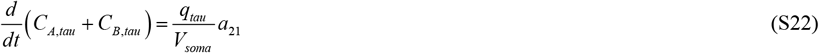

The numerical analysis of Eqs. (S19) and (S20) with initial conditions (11a,b) indicates that *C*_*A,tau*_ → 0 as *t* →∞. This enables the derivation of an approximate solution for *C*_*B, Aβ*_ that is applicable for large values of *t*:

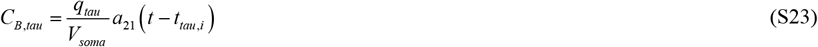

Also, substituting Eq. (S23) into Eq. (S20) yields:

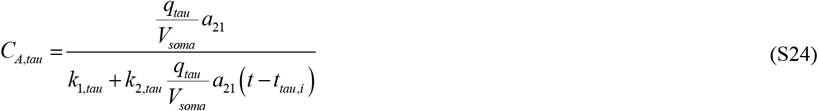

From Eqs. (S21) and (11c) it can be deduced that:

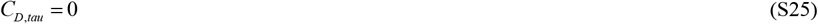

Employing Eqs. (15) and (S23) produces

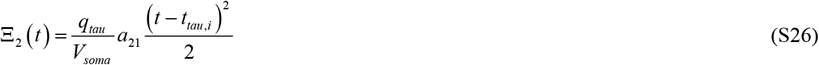

It is important to note that Eq. (S26) is derived under the assumption that *θ*_1/2,*B,tau*_ →∞. This differs from the conditions used to compute Figs. 3, S3, and S4, which were computed for the case of *θ*_1/2,*B,tau*_ →10^7^ s (refer to Table 2).

#### S2.2. The scenario with a fast deposition rate of tau aggregates into NFTs, *θ*_1/2,*B,tau*_ →0, as well as *T*_1/ 2, *A,tau*_ →∞, *T*_1/ 2, *B,tau*_ →∞, and *T*_1/ 2, *D,tau*_ →∞

In the case of immediate deposition of tau aggregates into NFTs, represented by *θ*_1/2,*B,tau*_ →0, Eqs. (8)–(10) simplify to:

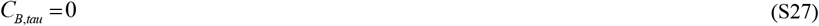

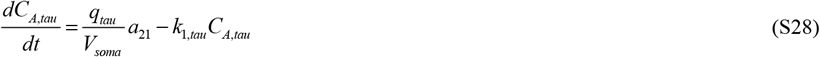

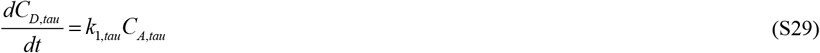

In this scenario, Eqs. (15) and (S27) result in:

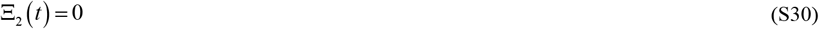

Adding Eqs. (S28) and (S29) yields:

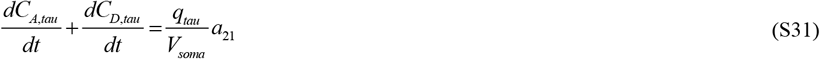

By integrating Eq. (S31) with respect to time and utilizing the initial conditions given by Eq. (11a,c), the following result is derived:

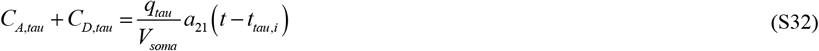

The increase in *C*_*A,tau*_ + *C*_*D,tau*_ over time is attributed to the synthesis of tau monomers in the soma, as well as the translocation of tau from the axonal to the somatodendritic compartments, a process typically observed in AD.

By substituting *C*_*A,tau*_ from Eq. (S32) into Eq. (S29), the following equation is obtained:

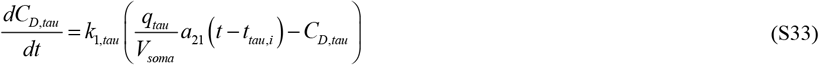

The exact solution of Eq. (S33), solved subject to initial condition (11c), is

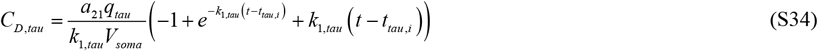

As *t* → 0, Eq. (S34) yields:

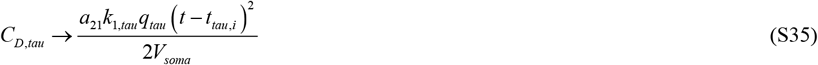

The quantity *C*_*A,tau*_ can be calculated by substituting Eq. (S34) into Eq. (S32) as follows:

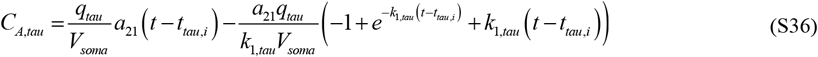

#### S3. A model for senile plaque growth

The methodology developed in ref. [49] serves as the foundation for this model. To simulate the growth of an Aβ plaque (Fig. 1), the approach focuses on calculating the total number of Aβ monomers, denoted as *N*_*Aβ*_, that have been incorporated into the plaque over time *t*. This calculation is based on the average concentration of Aβ monomers deposited into the plaque, following ref. [51]:

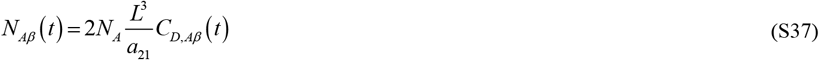

where the factor of 2 on the right-hand side of Eq. (37) accounts for the fact that the Aβ plaque develops between two neighboring neurons. Here, *N*_*A*_ represents Avogadro’s number. The membranes of individual neurons release Aβ monomers at a rate of *q*_*A*,0,*Aβ*_, contributing to the formation of the plaque (Fig. 1).

An alternative method, which also follows from ref. [51], involves calculating *N*_*Aβ*_ (*t*) based on the volume of a single Aβ plaque, *V*_*ABP*_ :

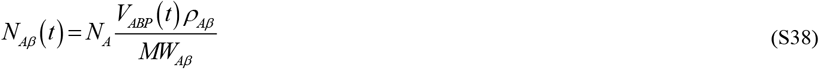

where *MW*_*Aβ*_ represents the average molecular weight of an Aβ monomer.

By equating the right-hand sides of Eqs. (S37) and (S38) and solving for the volume of the Aβ plaque, the following equation is derived:

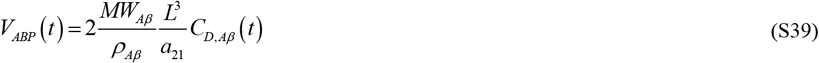

where 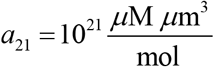 is the conversion factor from 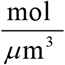 to *μ*M.

Assuming that the Aβ plaque is spherical,

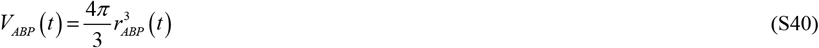

By equating the right-hand sides of Eqs. (S39) and (S40) and solving for the radius of the Aβ plaque, the following equation is obtained:

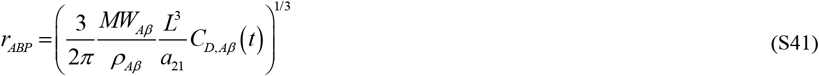

### S4. Numerical Solution

The numerical solutions of Eqs. (4)–(6) with initial conditions specified in Eq. (7), as well as of Eqs. (8)–(10) with initial conditions specified in Eq. (11), were obtained using MATLAB’s ODE45 solver (MATLAB R2024a, MathWorks, Natick, MA, USA). This solver is known for its accuracy and reliability. To achieve high precision in the results, both the relative and absolute tolerances, RelTol and AbsTol, were set to 1e-10.

### S5. Supplementary figures

**Fig. S1.**
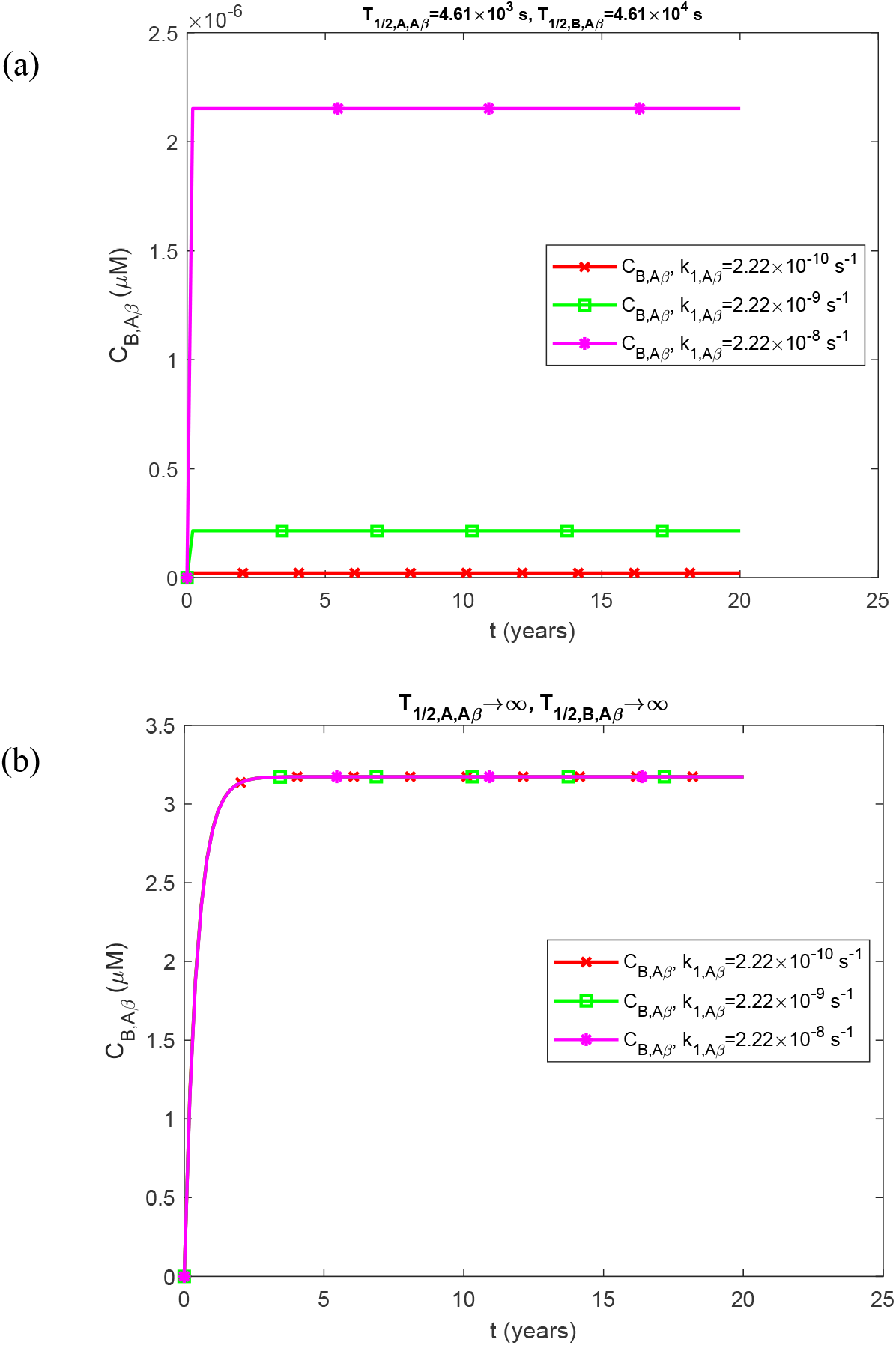
Similar to Fig. 2, but showing only the molar concentration of free Aβ aggregates (oligomers), *C*_*B, Aβ*_, with a larger *y*-axis scale. (a) This scenario represents physiologically relevant half-lives of Aβ monomers and aggregates. (b) This scenario illustrates dysfunctional protein degradation machinery corresponding to infinitely long half-lives of Aβ monomers and aggregates. Results are presented for three different values of nucleation rate constants for free Aβ aggregates, *k*_1,*Aβ*_. Parameters: *T*_1/2,*D,Aβ*_ →∞, *θ*_1/2,*B,Aβ*_ =10^7^ s.

**Fig. S2.**
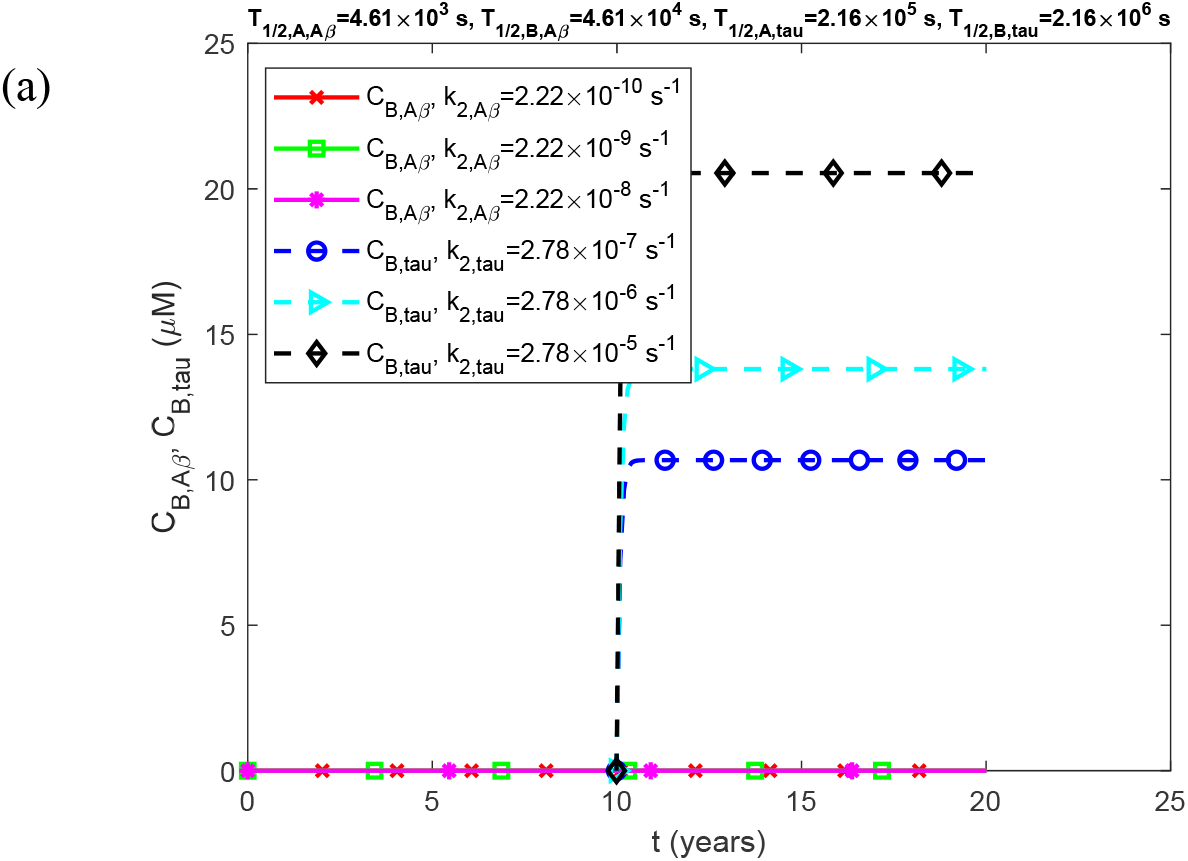
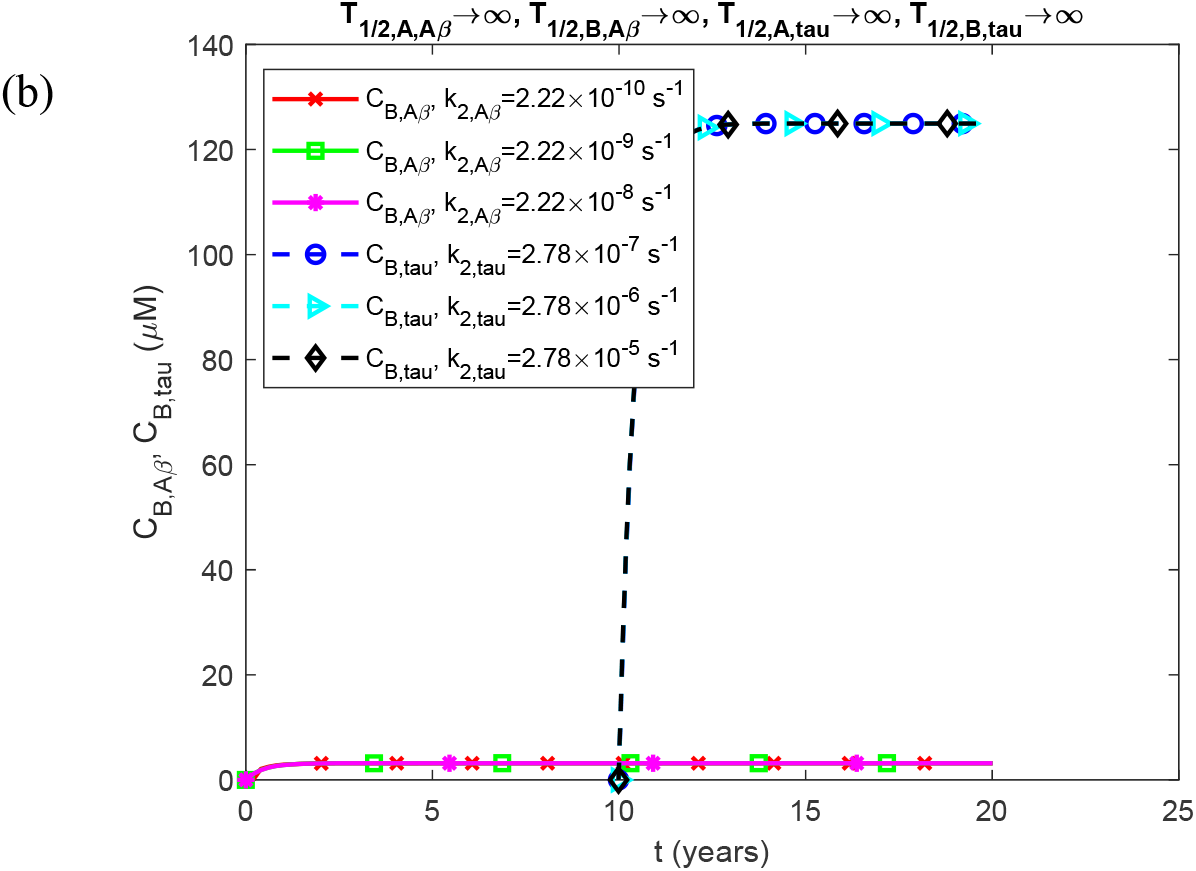
Molar concentrations of free Aβ aggregates (oligomers), *C*_*B, Aβ*_, and free tau aggregates (oligomers), *C*_*B,tau*_, as a function of time (in years). (a) This panel depicts the scenario with physiologically relevant half-lives for Aβ and tau monomers and aggregates. (b) This panel depicts the scenario involving dysfunctional protein degradation machinery, which corresponds to infinitely long half-lives of Aβ and tau monomers and aggregates. Results are plotted for three different values of the autocatalytic growth rate constants for free Aβ and tau aggregates, *k*_2, *Aβ*_ and *k*_2,*tau*_, respectively. Parameters: *T*_1/2,*D,Aβ*_ →∞, *T*_1/2,*D,tau*_ →∞, *θ*_1/2,*B,Aβ*_ =10^7^ s, and *θ*_1/2,*B,tau*_ =10^7^ s.

**Fig. S3.**
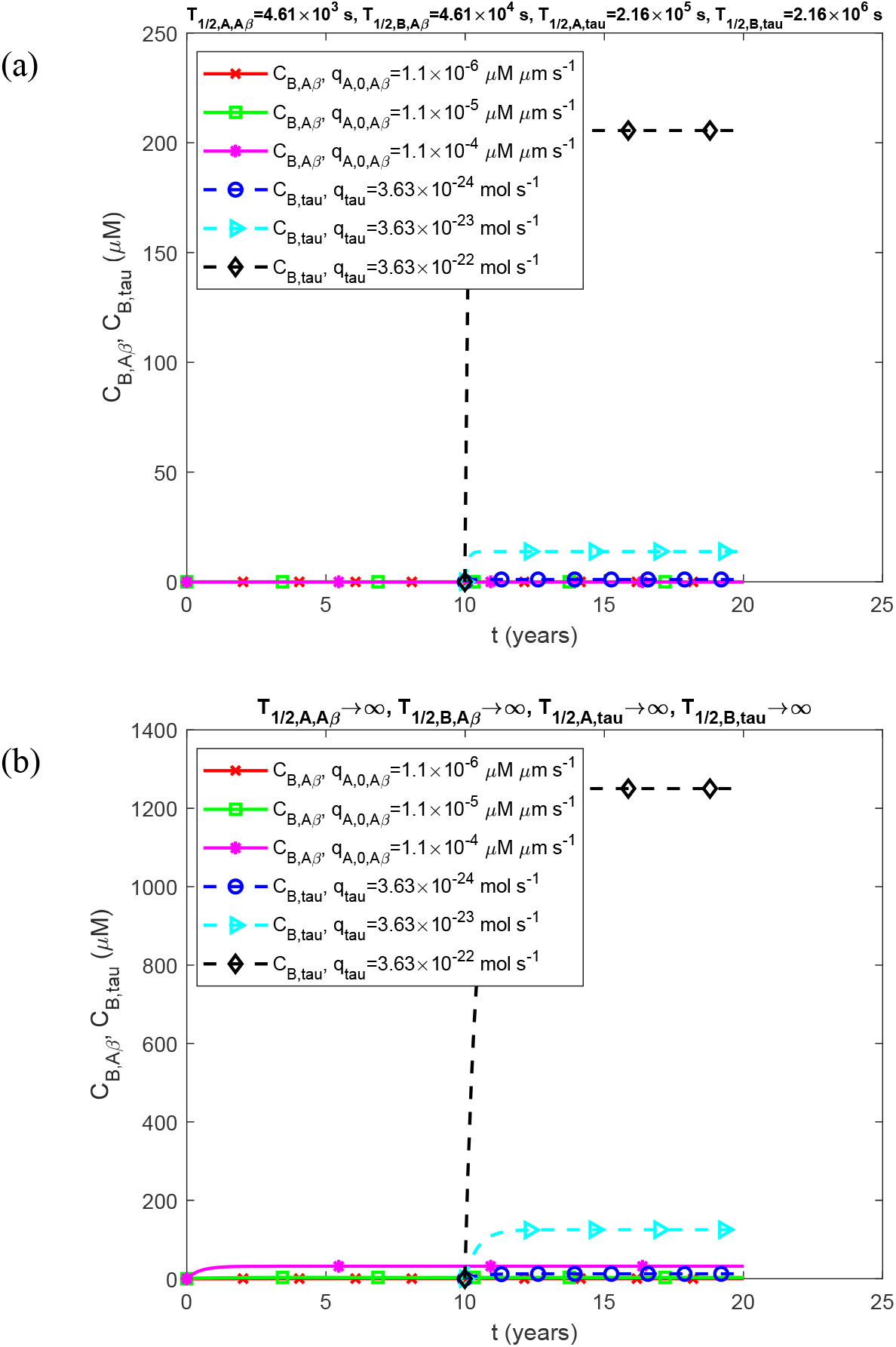
Molar concentrations of free Aβ aggregates (oligomers), *C*_*B, Aβ*_, and free tau aggregates (oligomers), *C*_*B,tau*_, as a function of time (in years). (a) This panel illustrates the scenario with physiologically relevant half-lives for Aβ and tau monomers and aggregates. (b) This panel depicts the scenario involving dysfunctional protein degradation machinery, corresponding to infinitely long half-lives of Aβ and tau monomers and aggregates. Results are presented for three different values of the production rates of Aβ and tau monomers, *q*_*A*,0, *Aβ*_ and *q*_*tau*_, respectively. Parameters: *T*_1/2,*D,Aβ*_ →∞, *T*_1/2,*D,tau*_ →∞, *θ*_1/2,*B,Aβ*_ =10^7^ s, and *θ* _1/2,*B,tau*_ =10^7^ s.

**Fig. S4.**
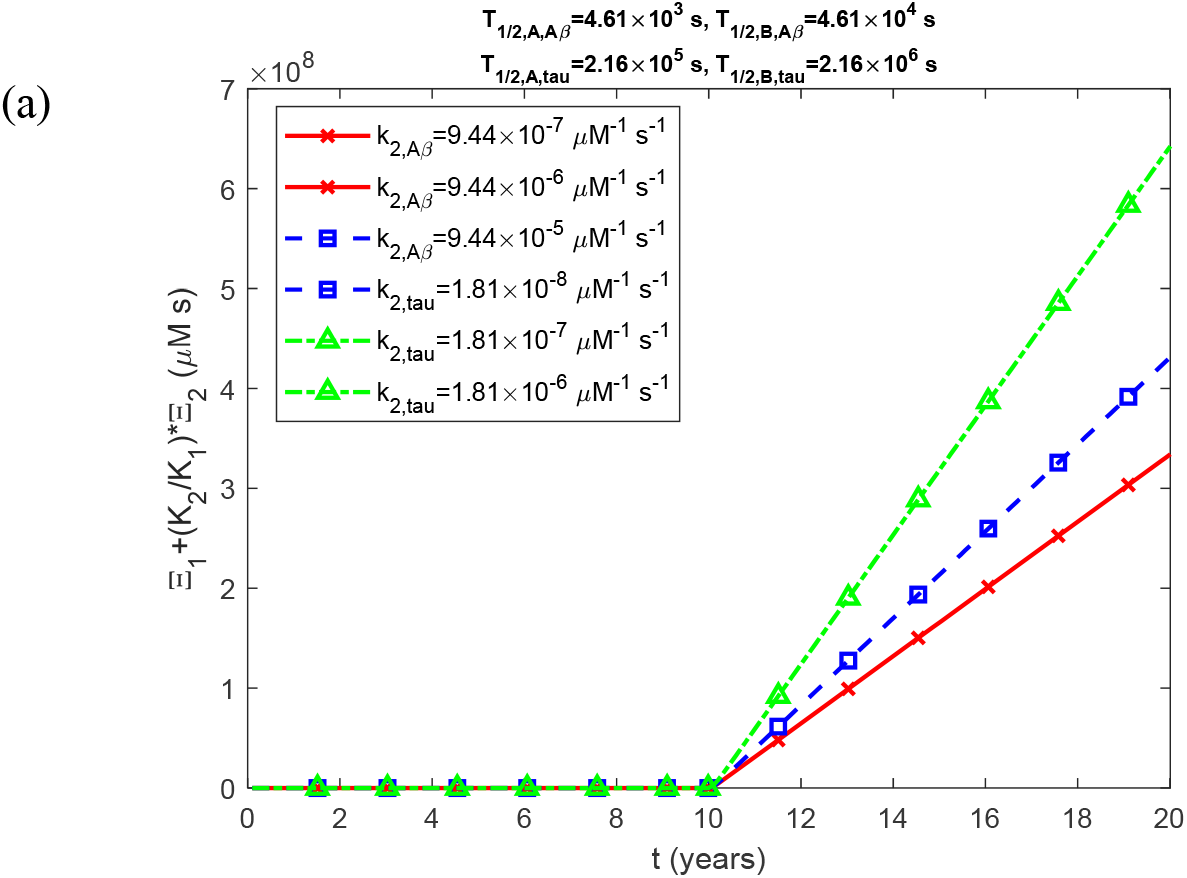
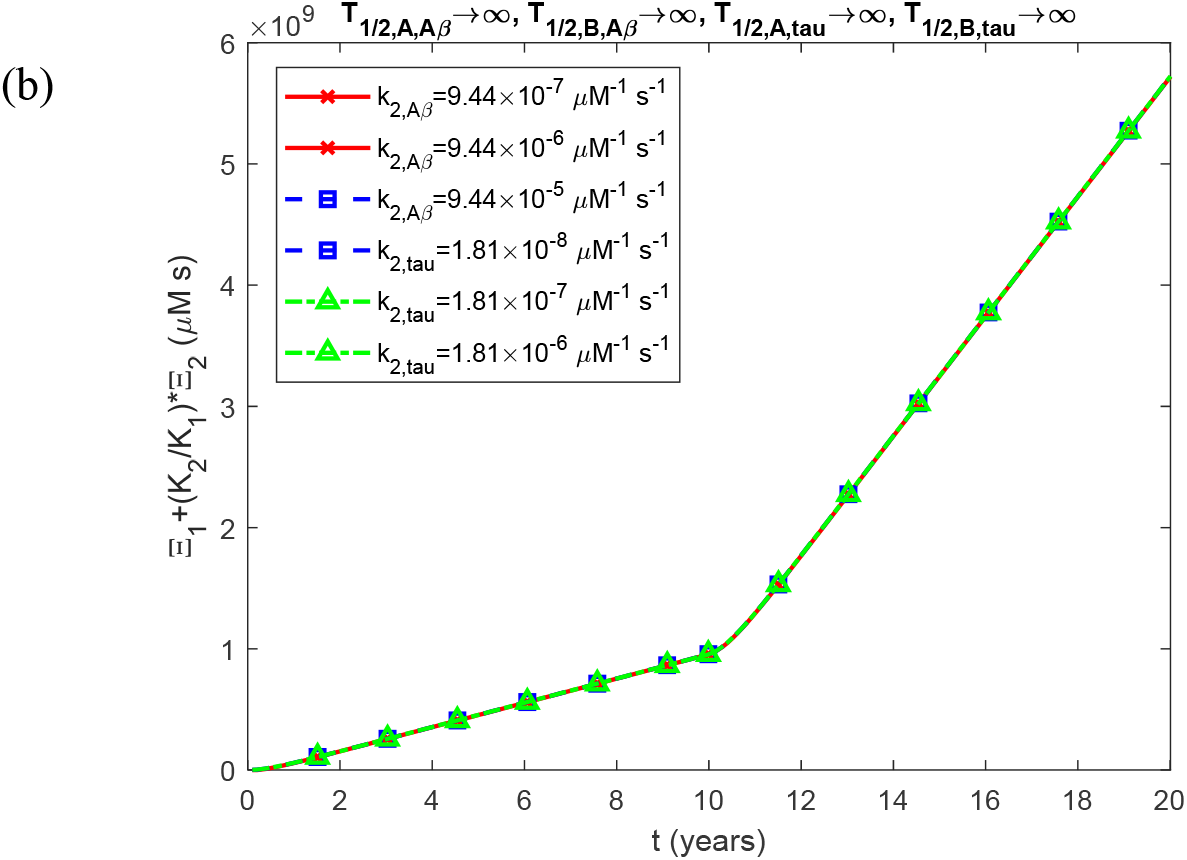
Combined accumulated toxicity of Aβ and tau oligomers, 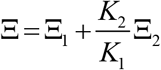, as a function of time (in years). (a) This panel shows results for physiologically relevant half-lives of Aβ and tau monomers and aggregates. (b) This panel represents the scenario with dysfunctional protein degradation machinery, corresponding to infinitely long half-lives of Aβ and tau monomers and aggregates. Results are presented for three different values of the autocatalytic growth rate constants for free Aβ and tau aggregates, *k*_2, *Aβ*_ and *k*_2,*tau*_, respectively. Parameters: *T*_1/2,*D,Aβ*_ →∞, *T*_1/2,*D,tau*_ →∞, *θ*_1/2,*B,Aβ*_ =10^7^ s, and *θ* _1/2,*B,tau*_=10^7^ s.

**Fig. S5.**
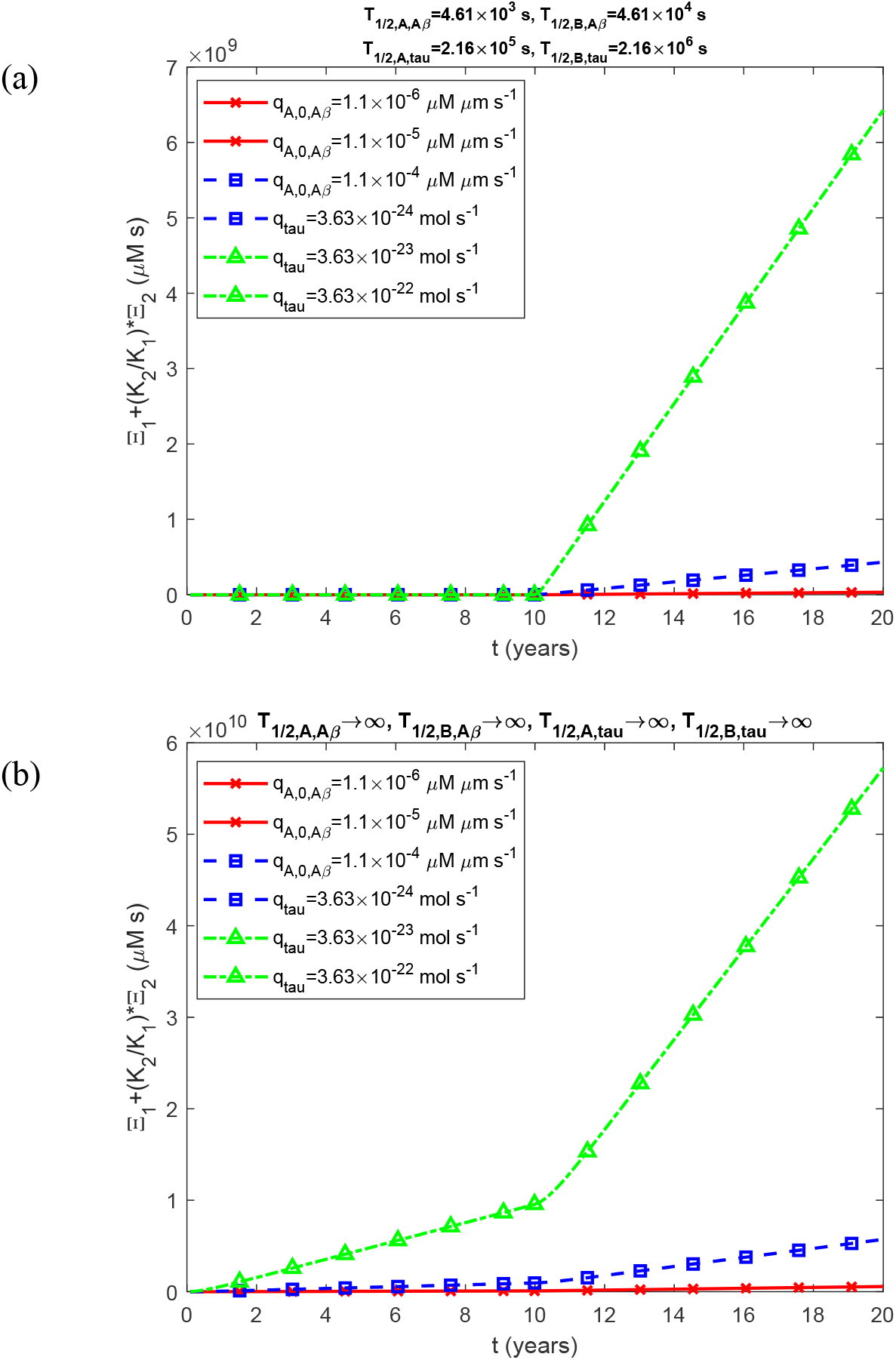
Combined accumulated toxicity of Aβ and tau oligomers, 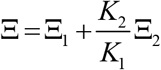, as a function of time (in years). (a) Results for the scenario with physiologically relevant half-lives of Aβ and tau monomers and aggregates. (b) Results for the scenario with dysfunctional protein degradation machinery, assuming infinitely long half-lives of Aβ and tau monomers and aggregates. Each panel presents results for three different values of production rates of Aβ and tau monomers, *q*_*A*,0, *Aβ*_ and *q*_*tau*_, respectively. Parameters: *T*_1/2,*D,Aβ*_ →∞, *T*_1/2,*D,tau*_ →∞, *θ*_1/2,*B,Aβ*_ =10^7^ s, and *θ* _1/2,*B,tau*_ =10^7^ s.

**Fig. S6.**
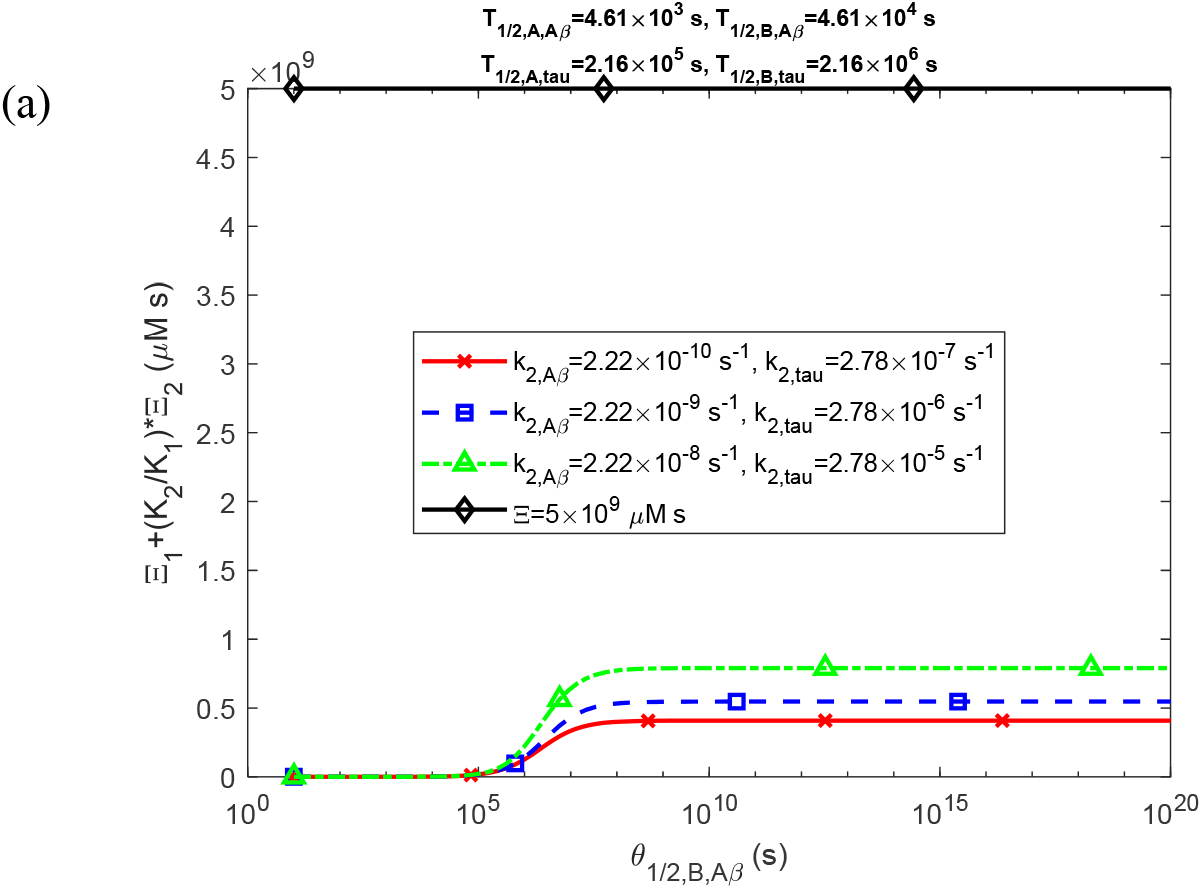
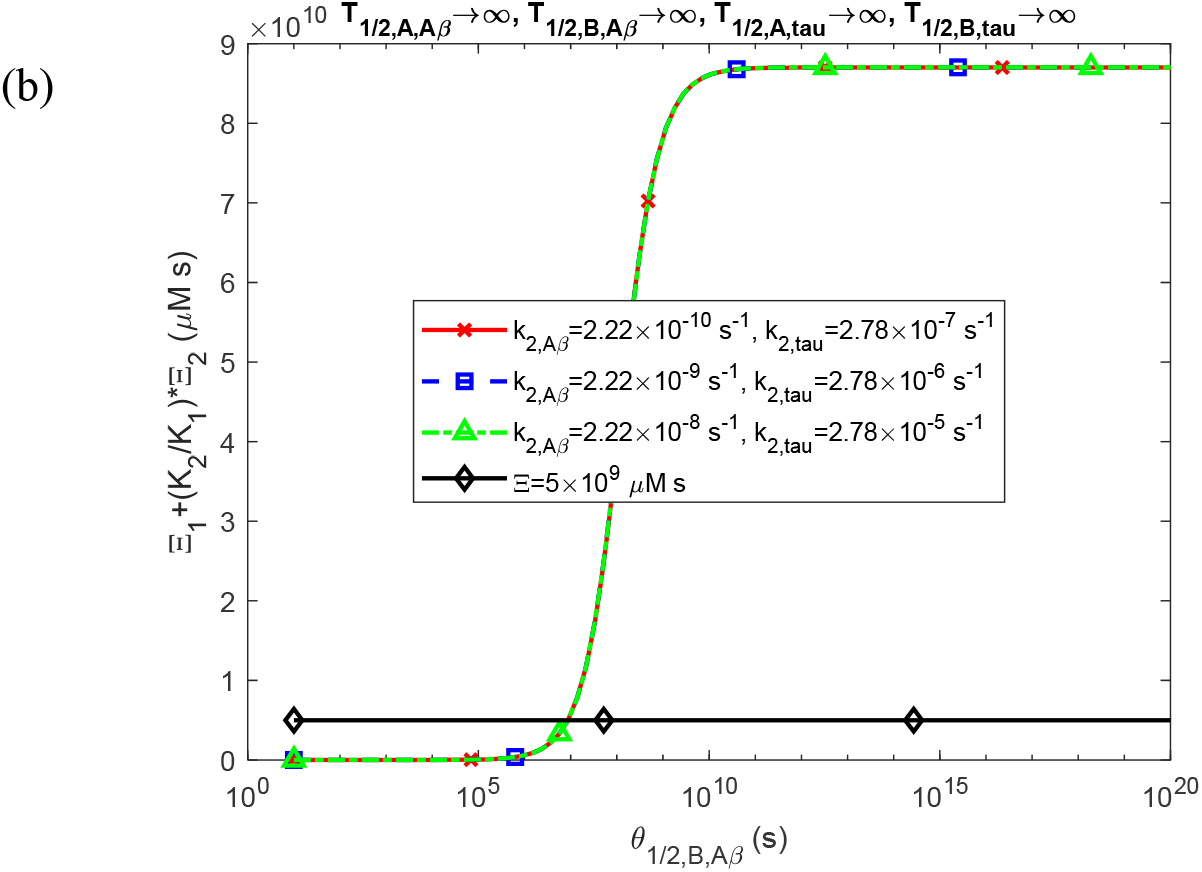
Combined accumulated toxicity of Aβ and tau oligomers, 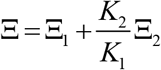, as a function of the half-deposition times, *θ*_1/2,*B,Aβ*_ and *θ*_1/2,*B,tau*_ (both set to equal values for this computation). (a) Results under the scenario with physiologically relevant half-lives of Aβ and tau monomers and aggregates. (b) Results for the scenario with dysfunctional protein degradation machinery, assuming infinitely long half-lives for both Aβ and tau monomers and aggregates. Data are shown at *t* = *t* _*f*_ = 6.31 × 10^8^ s for three different values of autocatalytic growth rate constants for free Aβ and tau aggregates, *k*_2, *Aβ*_ and *k*_2,*tau*_, respectively. Parameters: *T*_1/2,*D,Aβ*_ →∞, *T*_1/2,*D,tau*_ →∞.

**Fig. S7.**
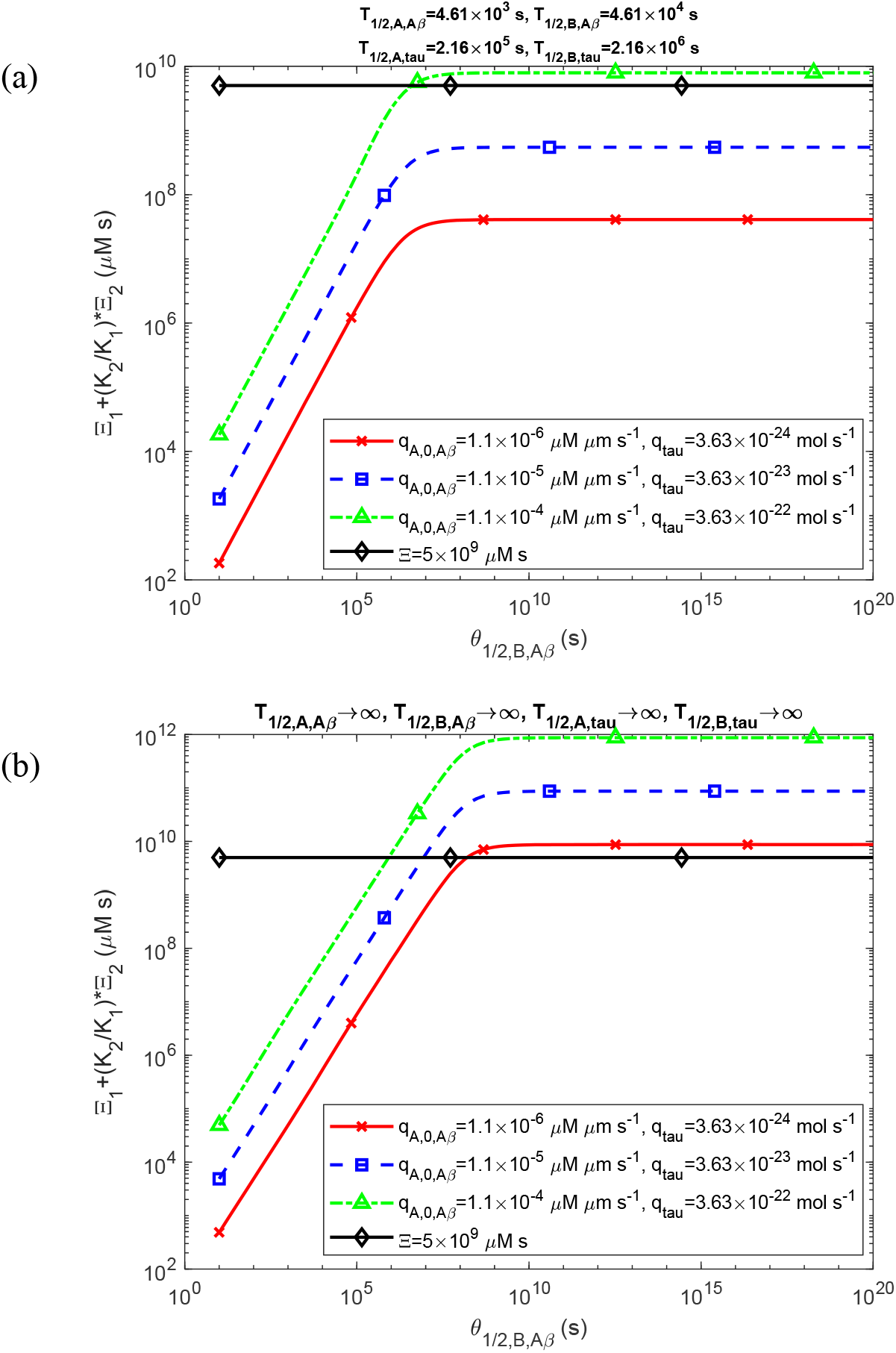
Combined accumulated toxicity of Aβ and tau oligomers, 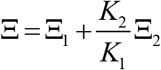, as a function of the half-deposition times, *θ*_1/2,*B,Aβ*_ and *θ*_1/2,*B,tau*_ (both set to equal values for this computation). (a) Results for the scenario with physiologically relevant half-lives for Aβ and tau monomers and aggregates. (b) Results for the scenario with dysfunctional protein degradation machinery, where Aβ and tau monomers and aggregates are assumed to have infinitely long half-lives. Data are shown at *t* = *t* _*f*_ = 6.31 × 10^8^ s for three different production rates of Aβ and tau monomers, *q*_*A*,0, *Aβ*_ and *q*_*tau*_, respectively. Parameters: *T*_1/2,*D,Aβ*_ →∞, *T*_1/2,*D,tau*_ →∞.

**Fig. S8.**
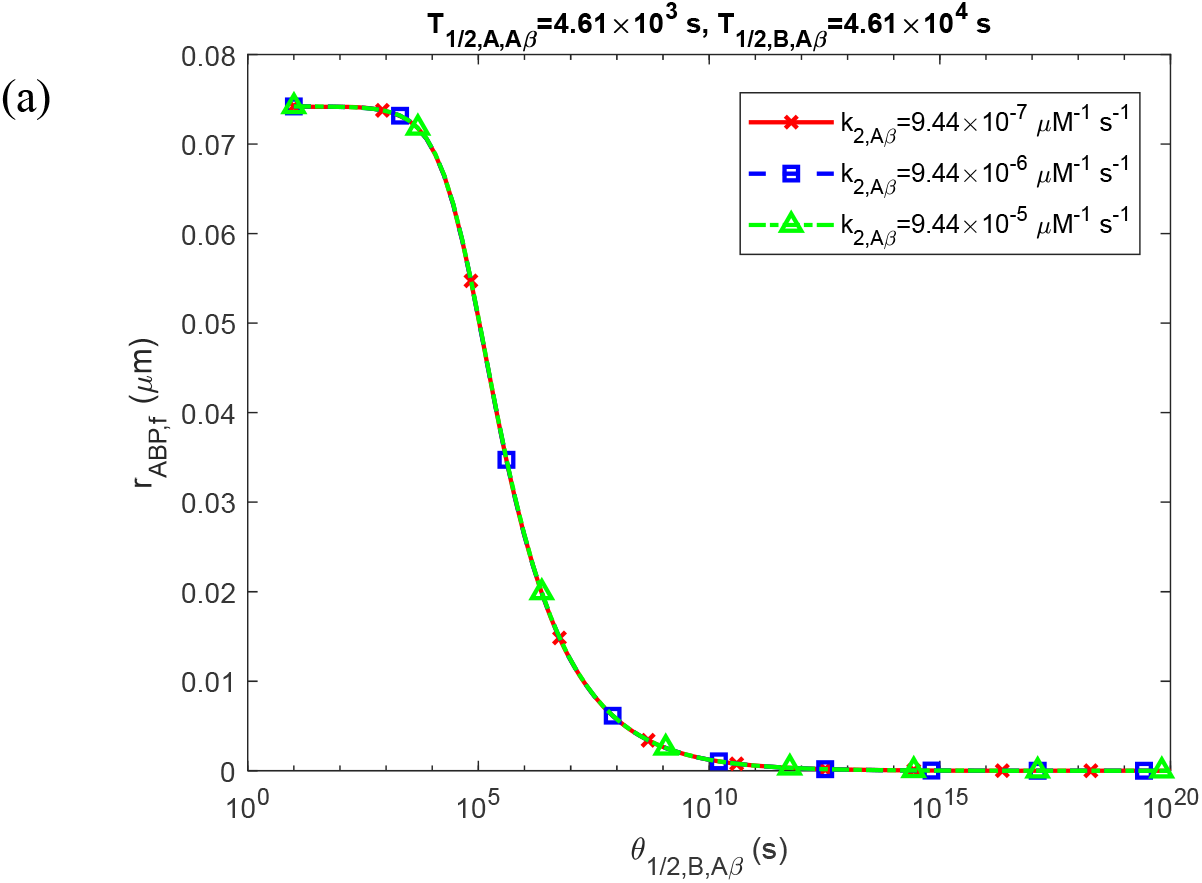
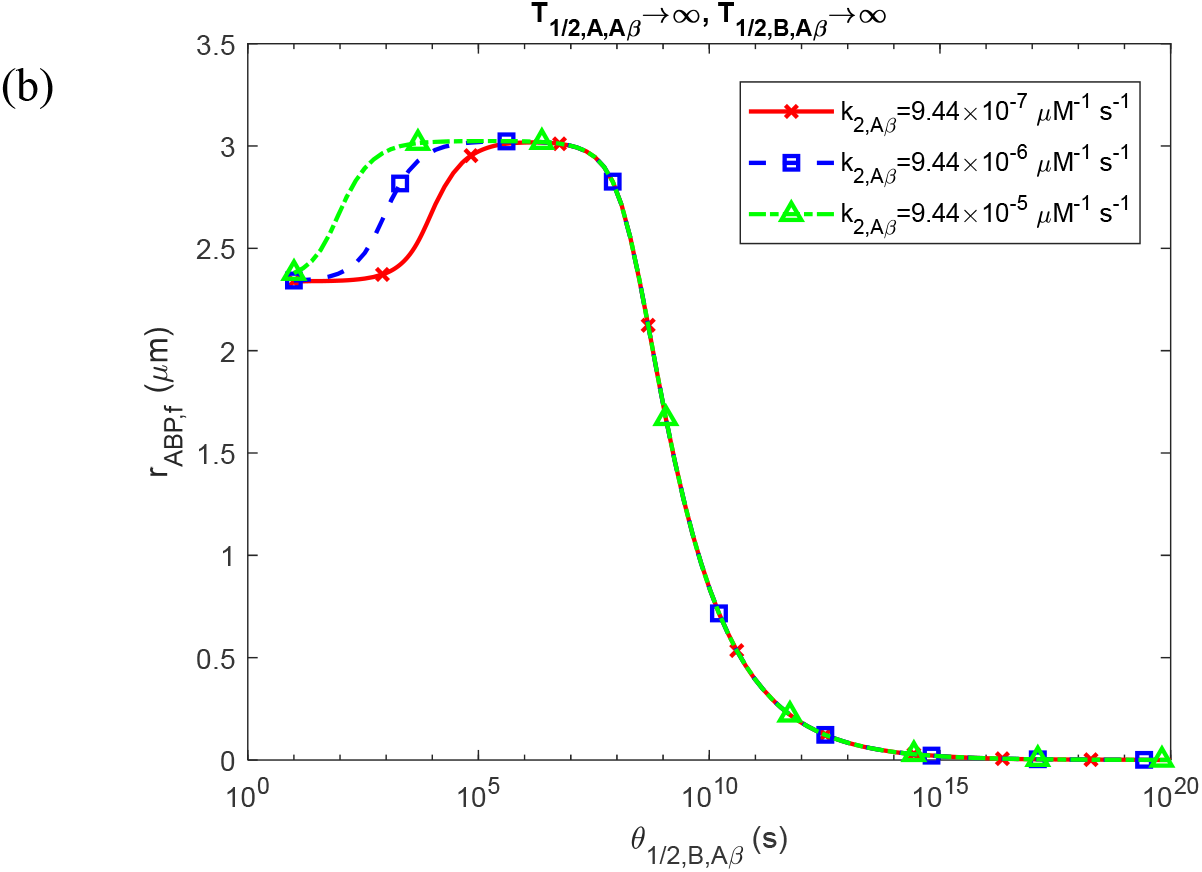
The radius of a growing Aβ plaque, *r*_*ABP, f*_, at *t* = *t* _*f*_ shown as a function of the half-deposition time *θ*_1/2,*B,Aβ*_. (a) This scenario represents physiologically relevant half-lives of Aβ monomers and aggregates. (b) This scenario depicts dysfunctional protein degradation machinery, characterized by infinitely long half-lives of Aβ monomers and aggregates. Results are presented for three different values of autocatalytic growth rate constants for free Aβ aggregates, *k*_2,*Aβ*_. Parameters: *t* _*f*_ = 6.31 × 10^8^ s, *T*_1/2,*D,Aβ*_ →∞.

**Fig. S9.**
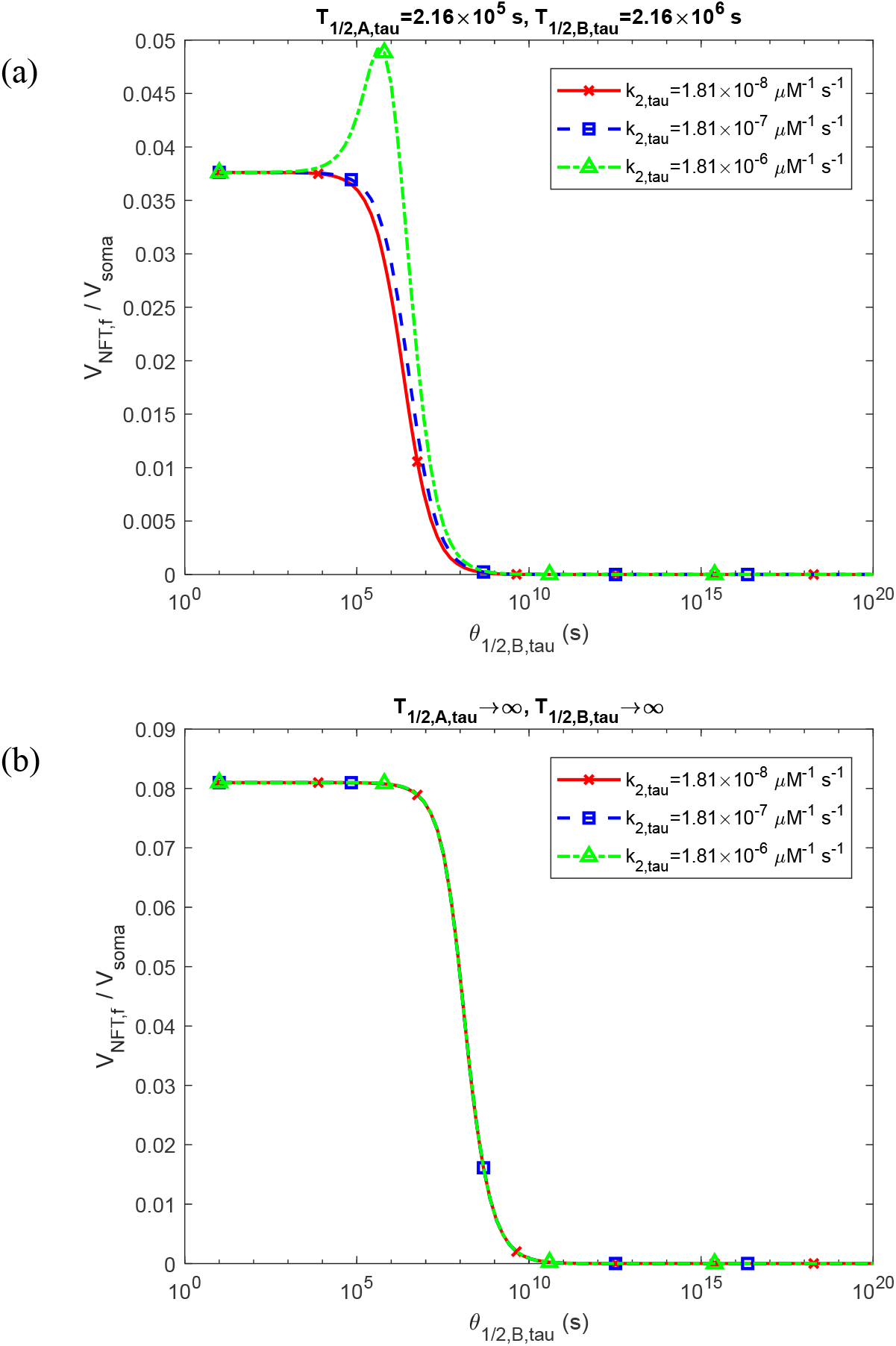
Fraction of the soma volume occupied by NFTs, 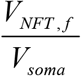, at *t* = *t* _*f*_ shown as a function of the half-deposition time *θ*_1/2,*B,Aβ*_. (a) Scenario representing physiologically relevant half-lives for tau monomers and aggregates. (b) Scenario illustrating dysfunctional protein degradation machinery, with infinitely long half-lives for tau monomers and aggregates. Results are presented for three different values of autocatalytic growth rate constants for free tau aggregates, *k*_2,*tau*_. Parameters: *t* _*f*_ = 6.31 × 10^8^ s, *T*_1/2,*D,tau*_ →∞.

**Fig. S10.**
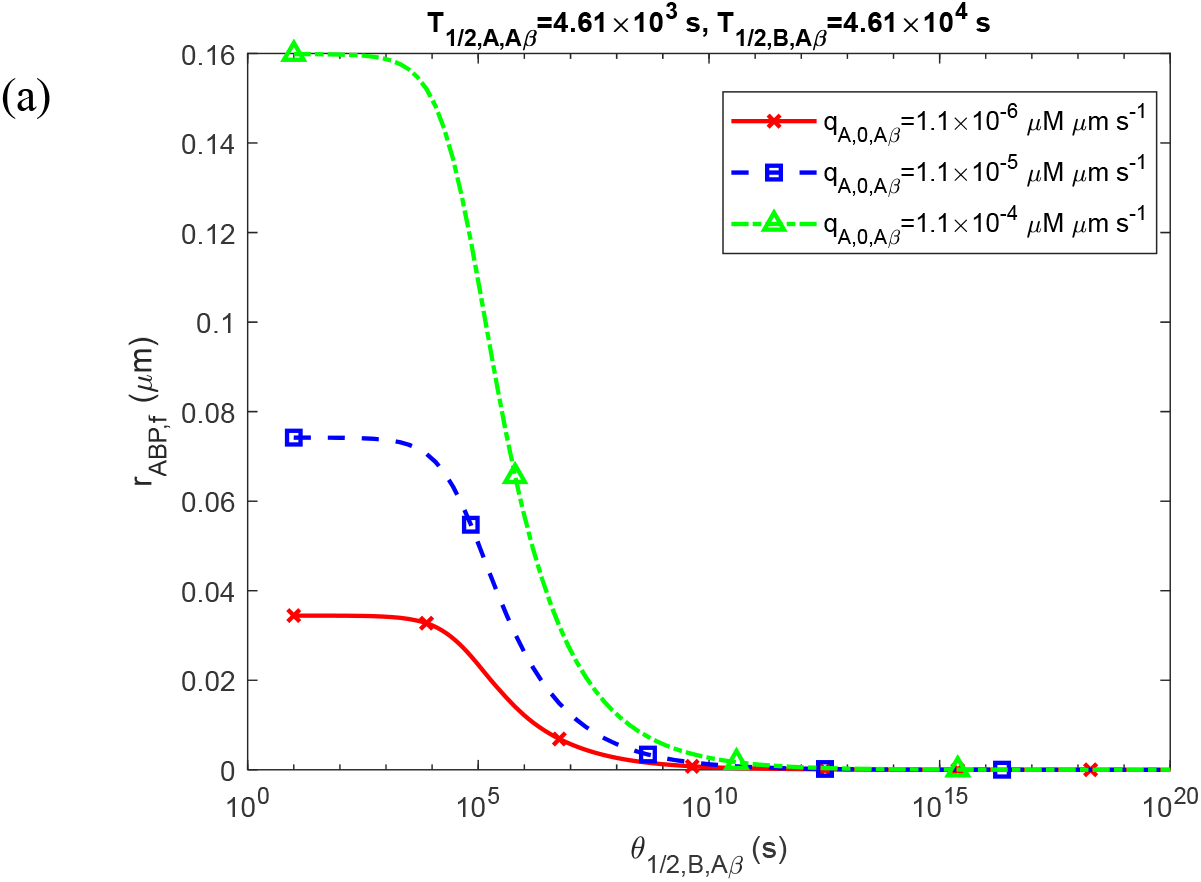
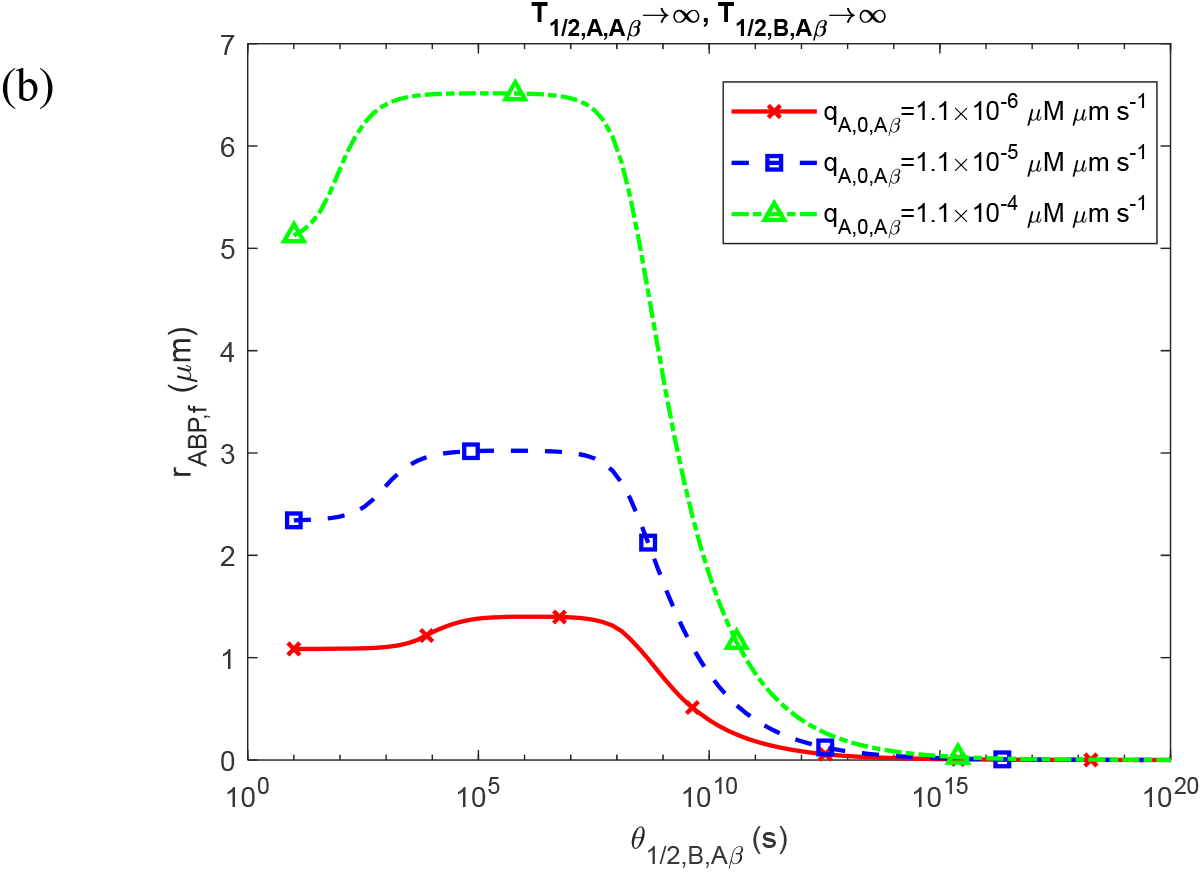
The radius of a growing Aβ plaque, *r*_*ABP, f*_, at *t* = *t* _*f*_ shown as a function of the half-deposition time *θ*_1/2,*B,Aβ*_. (a) This scenario represents physiologically relevant half-lives of Aβ monomers and aggregates. (b) This scenario depicts dysfunctional protein degradation machinery, characterized by infinitely long half-lives of Aβ monomers and aggregates. Results are presented for three different values of production rates of Aβ monomers, *q*_*A*,0,*Aβ*_, respectively. Parameters: *t* _*f*_ = 6.31 × 10^8^ s, *T*_1/2,*D,Aβ*_ →∞.

**Fig. S11.**
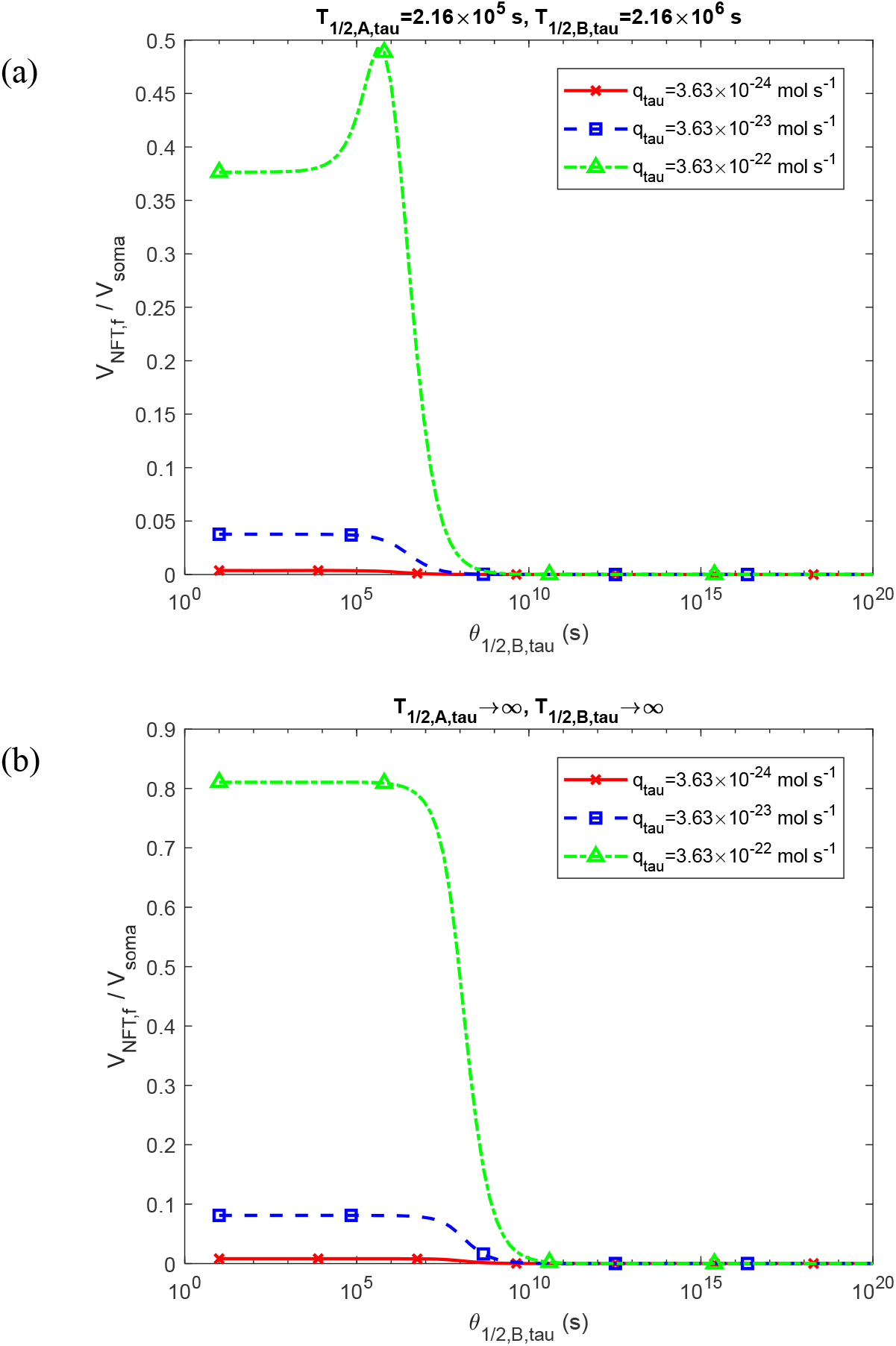
Fraction of the soma volume occupied by NFTs, 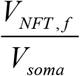, at *t* = *t* _*f*_ shown as a function of the half-deposition time *θ*_1/2,*B,Aβ*_. (a) Scenario representing physiologically relevant half-lives for tau monomers and aggregates. (b) Scenario illustrating dysfunctional protein degradation machinery, with infinitely long half-lives for tau monomers and aggregates. Results are presented for three different values of supply rates of tau monomers, *q*_*tau*_. Parameters: *t* _*f s*_= 6.31 × 10^8^ s, *T*_1/2,*D,tau*_ →∞.

